# Anaplerotic nutrient stress drives synergy of angiogenesis inhibitors with therapeutics targeting tumor metabolism

**DOI:** 10.1101/2023.05.07.539744

**Authors:** Sunada Khadka, Yu-Hsi Lin, Jeffrey Ackroyd, Yi-An Chen, Yanghui Sheng, Wubin Qian, Sheng Guo, Yining Chen, Eliot Behr, Yasaman Barekatain, Md. Nasir Uddin, Kenisha Arthur, Victoria Yan, Wen-Hao Hsu, Qing Chang, Anton Poral, Theresa Tran, Surendra Chaurasia, Dimitra K. Georgiou, John M. Asara, Floris P. Barthel, Steve W. Millward, Ronald A. DePinho, Florian L. Muller

## Abstract

Tumor angiogenesis is a cancer hallmark, and its therapeutic inhibition has provided meaningful, albeit limited, clinical benefit. While anti-angiogenesis inhibitors deprive the tumor of oxygen and essential nutrients, cancer cells activate metabolic adaptations to diminish therapeutic response. Despite these adaptations, angiogenesis inhibition incurs extensive metabolic stress, prompting us to consider such metabolic stress as an *induced vulnerability* to therapies targeting cancer metabolism. Metabolomic profiling of angiogenesis-inhibited intracranial xenografts showed universal decrease in tricarboxylic acid cycle intermediates, corroborating a state of anaplerotic nutrient deficit or stress. Accordingly, we show strong synergy between angiogenesis inhibitors (Avastin, Tivozanib) and inhibitors of glycolysis or oxidative phosphorylation through exacerbation of anaplerotic nutrient stress in intracranial orthotopic xenografted gliomas. Our findings were recapitulated in GBM xenografts that do not have genetically predisposed metabolic vulnerabilities at baseline. Thus, our findings cement the central importance of the tricarboxylic acid cycle as the nexus of metabolic vulnerabilities and suggest clinical path hypothesis combining angiogenesis inhibitors with pharmacological cancer interventions targeting tumor metabolism for GBM tumors.

## INTRODUCTION

Angiogenesis inhibitors, particularly Avastin, a humanized, anti-hVEGFA monoclonal antibody, have become a mainstay for the treatment of vascularized solid tumors as a first line or second line therapy^1^. Avastin significantly restricts blood flow to the tumor and given in combination with chemotherapy, can elicit a survival benefit in some cancers^2,3^. However, the tumor static effects of angiogenesis inhibition are often transient due to a multitude of cancer cell intrinsic and extrinsic adaptive mechanisms^4–6^. Despite this limitation, anti-angiogenic therapies such as Avastin still remain important in the treatment of many cancer types, including glioblastoma multiforme (GBM), which is a highly vascularized, infiltrative, and invariably fatal disease^7^. In GBM patients, Avastin alone or in combination with chemotherapy, alleviates cancer-related symptoms, elicits a radiological response, and improves progression-free survival^7,8^. However, a transient response combined with the invasiveness of glioma cells and their co-option of existing vasculature has dampened the clinical utility of Avastin in GBM^7^. At the same time, angiogenesis inhibition can instigate significant tumor-intrinsic and -extrinsic metabolic adaptations to sustain tumor growth, prompting speculation of targetable induced vulnerabilities^9,10^.

Preclinical studies have demonstrated that metabolic rewiring by the tumors is a proximal consequence of angiogenesis inhibition^3,5,11^. In support, clinical magnetic resonance imaging (MRI) studies show reduced tumor perfusion^12–14^ and subsequent induction of intratumoral hypoxia in response to anti-angiogenic treatments^2,13,15–17^. One prominent response to angiogenesis inhibition-induced oxygen deficiency is HIF-1α stabilization, which can trigger a cascade of metabolic adaptations to facilitate tumor growth and metastasis in an oxygen- and nutrient-deficient microenvironment^18,19^. The elucidation of such adaptive mechanisms could reveal cancer-specific vulnerabilities that may inform more effective combination therapies.

Glucose and glutamine are among the most critical nutrients that tumors derive from blood^20^. Tumors consume glucose voraciously to generate biosynthetic precursors necessary for proliferation via glycolysis; this includes ribose (glucose-6-phosphate—pentose phosphate pathway), purines (3-phosphoglycerate—one carbon metabolism), and lipid head groups (dihyroxyacetone-phosphate—glycerol)^20,21^. While pyruvate, the end-product of glycolysis, is mostly secreted as lactate to enable NAD+ regeneration and maintain glycolytic flux, a significant portion of glucose-derived pyruvate also enters the tricarboxylic acid (TCA) cycle^20,21^. Glutamine also converges on the TCA cycle and serves as a carbon and nitrogen donor for nucleotide synthesis and transamination reactions and is essential for glutathione generation^20,22^. Continued proliferation of cancer cells is contingent upon continuous replenishment of TCA cycle carbon atoms (anaplerosis) that are drained for macromolecule biosynthesis (cataplerosis)^23^. Depletion of TCA cycle intermediates is characteristic of Avastin-treated tumors, which suggests that angiogenesis inhibition induces anaplerotic nutrient stress^24^.

To better understand how angiogenesis inhibition influences tumor metabolism and might inform on effective combination therapies, we characterized the metabolomic and transcriptomic profile of intracranial orthotopic gliomas on Avastin treatment. Angiogenesis inhibition led to a marked reduction in TCA-cycle metabolites, indicative of anaplerotic nutrient stress. Indeed, we demonstrate robust anti-tumor synergy and profound nutrient stress with combined Avastin and glycolysis/OxPhos inhibitor treatment. Our findings indicate that disruptions to energy metabolism in tumors treated with angiogenesis inhibitors constitute an “induced vulnerability” that invite a systematic study of how such vulnerabilities might be exploited to potentiate the efficacy of angiogenesis inhibition.

## RESULTS

### Angiogenesis inhibition enhances hypoxia and anaplerotic nutrient stress in intracranial tumors

We implanted intracranial *ENO1*-deleted GBM xenografts into *Foxn1^nu/nu^* nude mice and administered the anti-human VEGFA antibody Avastin – which neutralizes (human) glioma cell-secreted human VEGFA but does not neutralize endogenous mouse VEGFA (**Supplementary Figure 1a**). After one week, we assessed the effect of the treatment on tumor growth with T1-weighted MRI with and without the non-permeable contrast enhancement agent gadobutrol (Gadavist ®) and T2-weighted MRI (**Figure 1a**). Untreated intracranial tumors exhibited edema, which was visible as a hyperintense region on T2-weighted MRI and on contrast-enhanced T1-weighted MRI (**Figure 1a**). In contrast, Avastin-treated tumors showed attenuated contrast enhancement, indicating restoration of mature tumor vasculature (normalized) and re-sealing of the BBB (**Figure 1a**) in a manner consistent with human tumor studies^25–30^. Thus, our findings confirm that intracranial D423 *ENO1*-deleted tumor xenografts have a breached BBB that can be resealed with angiogenesis inhibitor treatment.

**Figure 1:**
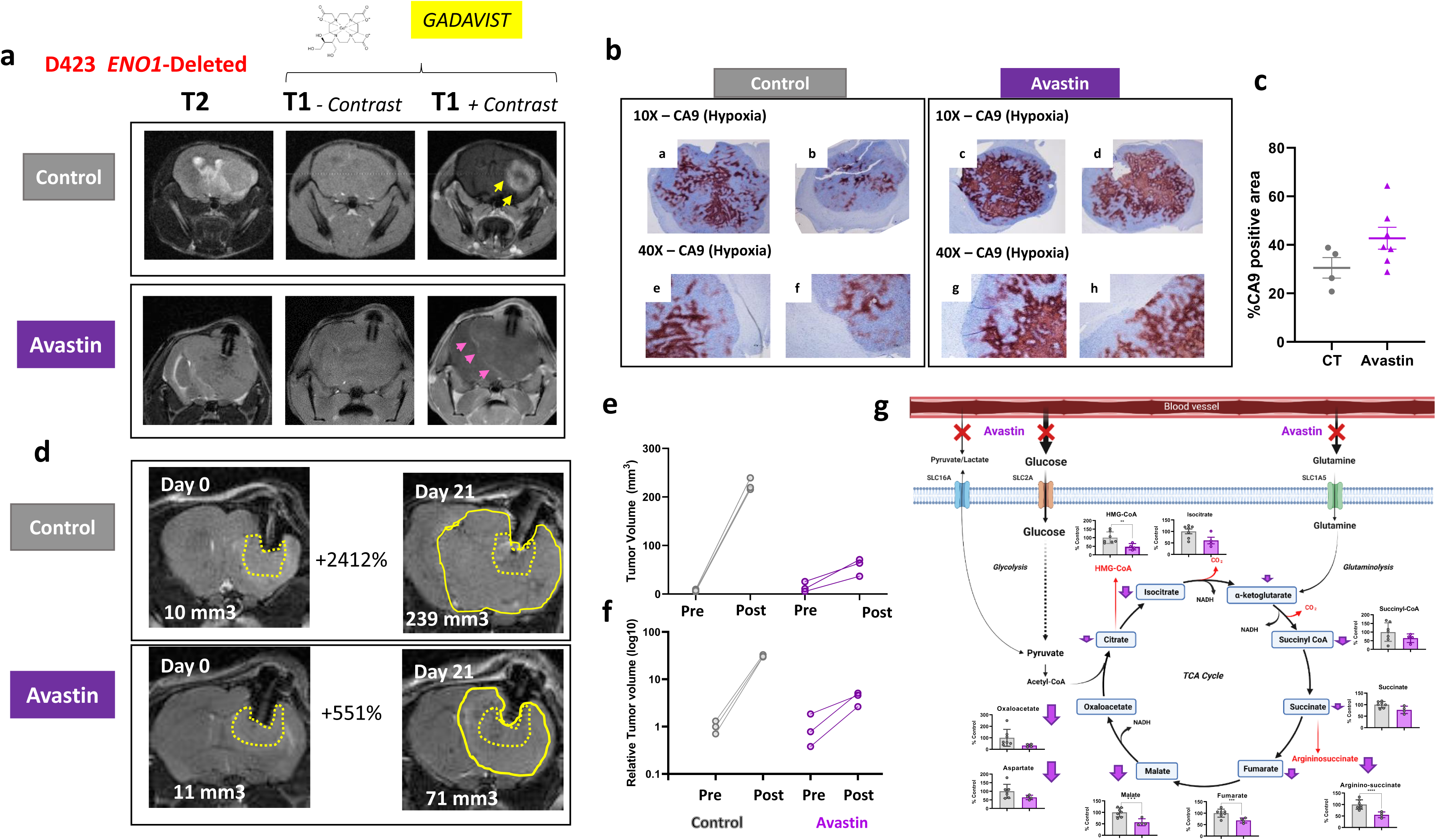
Angiogenesis inhibition re-seals the breached blood brain barrier, impairs perfusion and nutrient import, and modestly inhibits tumor growth. **a.** Intracranial tumors were generated by implanting D423 *ENO1* homozygously deleted glioma cells in Foxn1*^nu/nu^* nude mice. Tumor growth was followed by T2-weighted MRI every two weeks. Animals in the Avastin group were treated with Avastin (5mpk, x2 per week). After one week of treatment, T1 MRI with and without contrast (Gadavist, IV) was performed on untreated and Avastin treated mice. Untreated tumors exhibited consistent contrast enhancement (yellow arrows), indicative of a breached blood-brain-barrier(BBB) in these tumors. Avastin treated tumors showed a dramatic decrease in contrast enhancement, with a residual weak signal apparent at the tumor edges (pink arrows), suggesting an effective resealing of the BBB by Avastin treatment. **b-c.** Avastin treatment causes significant elevation in hypoxia, determined by the carbonic anhydrase 9 (CA9, a hypoxia marker) staining in intracranial tumors (**b**). Quantification of hypoxic areas (brown) in tumors are plotted (**c**). **d-e.** T2-weighted MRI scans showing the size of the tumors on day zero (dotted yellow line) and on day 21 (solid yellow line). Tumor volumes are indicated on the bottom left corner of each image. Pre- and post-treatment tumor volume comparisons (raw and relative) of control and Avastin treated mice after 21 days of treatment. **f.** Angiogenesis inhibition by Avastin significantly impairs import of major blood borne anaplerotic nutrients into the tumors, resulting in significant metabolic stress in tumors. Avastin treatment leads to a global decrease in TCA cycle intermediates, which are crucial for the generation of biosynthetic intermediates in cancer cells. Data are mean ± SD. Asterik(*) represents statistical significance (p<0.05) achieved by two-tailed t-test (**g**).

While vascular normalization improves perfusion, multiple studies have documented elevated tumor hypoxia as a proximal consequence of angiogenesis inhibition^25^(**Supplementary Figure 2a-b**). Specifically, the hypoxia imaging PET tracer ^18^F-fluoromisonidazole (FMISO) shows increased retention following Avastin treatment^31^. Consistent with these clinical observations, post-mortem histopathological analyses of Avastin-treated gliomas revealed a substantial increase in hypoxic areas, as evidenced by elevated CA9 expression (**Figure 1b-c**) as well as impaired tumor growth but no regressions (**Figure 1d-f**). These observations prompted us to consider that angiogenesis inhibition may also reduce access to blood-borne nutrients needed to meet the anabolic demands of rapidly proliferating cancer cells (**Supplementary Figure 1c-e**). To directly test this notion, we conducted an unbiased metabolomics analysis of intracranial tumors treated with Avastin, which revealed a substantial reduction in the levels of TCA cycle intermediates, including oxaloacetate, citrate, and fumarate (**Figure 1g**). Together, these results indicate that Avastin treatment spurs metabolic stress in intracranial tumors.

### Re-sealing of the blood-brain barrier does not abrogate glycolysis inhibition nor the anti-neoplastic effect of the phosphonate enolase inhibitor HEX

We previously reported the anti-neoplastic efficacy of a phosphonate inhibitor of enolase (“HEX”) in an intracranial orthotopic murine model of GBM^32^. As resealing the BBB is a hallmark of tumors treated with angiogenesis inhibitors^25–30^ and HEX shows poor BBB penetration^33^, we employed T2-weighted MRI to assess tumor growth of Avastin and HEX treatments, alone and in combination (**Figure 2a**). Treatment with Avastin or HEX slowed the growth of *ENO1*-deleted gliomas and was not associated with frank tumor regression (**Figure 2a-d**); whereas combined treatment resulted in profound tumor regression (**Figure 2a-d**) and increased survival: 67 days for combined therapy compared with 42 days for HEX, 48 days for Avastin (median: 48 days), and 31 days for vehicle control (**Figure 2e**), without any adverse effects (**Supplementary Figure 3a-d**). Consistent with therapeutic synergy, tumors showed markedly decreased proliferation (phosphor-histone H3) and increased apoptosis (cleaved caspase 3) relative to monotherapy and control tumors (**Figure 2f-h**).

**Figure 2:**
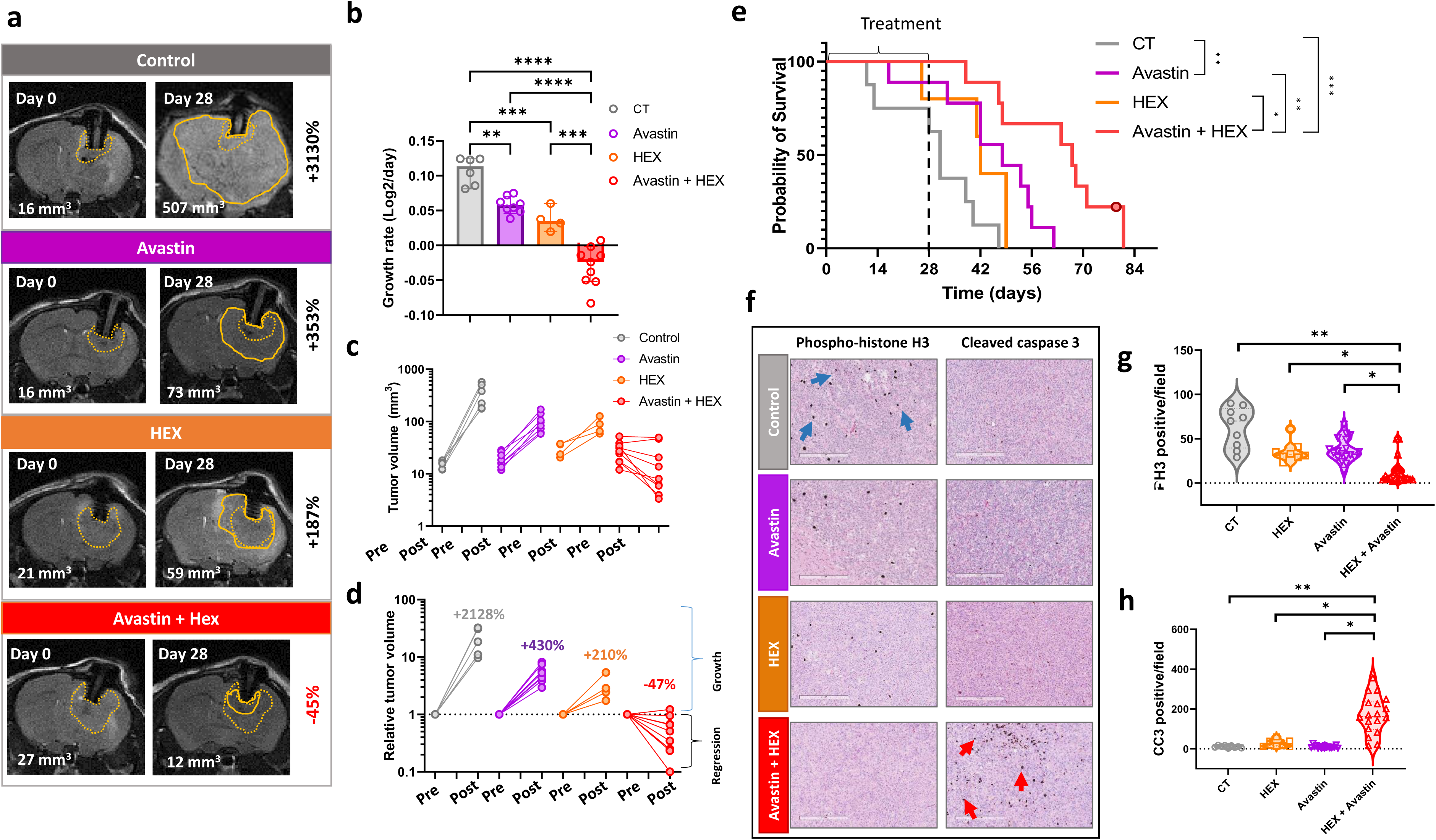
Long-term combination of angiogenesis inhibitor Avastin and Enolase inhibitor HEX results in regression of intracranial tumors at doses where either drug is only tumor-static as a monotherapy. Intracranial tumors were generated in Foxn1 *^nu/nu^* nude mice by implanting D423 *ENO1* homozygously deleted glioma cells and tumor growth was monitored every week by T2-weighted MRI. Four weeks later, when tumors reached approximately 20 mm^3^ in volume, mice were separated into four groups; control(vehicle), Avastin (5 mpk, 2X per week), Enolase inhibitor HEX (225 mpk, 12X per week) or Avastin plus HEX (Avastin, 5mpk 2X per week + 225 mpk SC 12X per week) were administered for 28 days. MRI scans were taken every two weeks to monitor tumor growth in response to treatment, and tumor volume changes were calculated. **a.** T2-weighted MRI images of animals before and after 28 days of treatment with tumor volumes indicated in mm^3^ in the lower left corner of the image; initial tumor outlines are shown in dotted yellow lines, while tumors after 28 days are shown in solid lines. **b-d**. Growth rates of tumors (**b**) and pre- and post-treatment tumor volume (**c-d**) in different treatment groups. Treatment with Avastin or HEX as single agents substantially attenuated tumor growth but did not result in actual tumor regression. However, the combination of Avastin with HEX resulted in tumor regression in all treated animals, and a complete eradication of tumors in some animals. Animals were taken off the treatments on day 28 and probability of survival after treatment discontinuation in each group was determined. Highlighted data point indicates mouse that died of reasons unrelated to tumor burden. **e**. Avastin and HEX combination lead to a significant extension of survival compared to HEX and Avastin alone. **f-i.** Histopathological analyses of brain tumor sections extracted from the mice show a dramatic reduction in phospho-histone H3 (PH3) positive cells (**f,g**) (an index of proliferation, blue arrows in the picture) in tumors treated with the combination of Avastin and HEX, concomitant to a dramatic increase in dying cells cleaved caspase 3 (CC3) positive cells (**f,h**), (an index of apoptosis, red arrows in picture), compared to the control, Avastin and HEX groups. Data are mean ± SD. Asterik(*) represents statistical significance (p<0.05) achieved by ordinary one-way ANOVA and Tukey’s multiple comparisons test (**b**,**g,h**).

To determine whether the accentuated anti-tumor activity of the combination of Avastin with HEX could be explained by enhanced glycolysis inhibition, we performed unbiased comprehensive metabolomic profiling of the single and combination treated intracranial tumors. Inhibition of glycolysis, evidenced by accumulation of glycolytic intermediates upstream of enolase reaction, was specifically observed only in tumors of mice treated with either HEX as a monotherapy or in combination with Avastin (**Supplementary Figure 3a**). The extent of disruption of glycolysis was comparable between tumors treated with either HEX as a monotherapy or in combination with Avastin (**Supplementary Figure 3a**).

Avastin only neutralizes human VEGFA secreted by human malignant xenografted cancer cells and does not bind to endogenous mouse Vegfa^34^. Murine stroma-secreted Vegfs could continue to promote neovascularization despite Avastin treatment, prompting us to pursue an orthogonal approach with Tivozanib, which is effective against both mouse and human VEGFR1/2/3^35^, either as a single agent or in combination with HEX (**Supplemental Figure 1b and 5a-e**). Tivozanib monotherapy yielded modest stasis of tumor growth, while combination therapy with HEX yielded significant regression of intracranial tumors, including complete tumor eradication (**Figure 2a-d and Supplemental Figure 5a-d**) and prolonged survival (63 days) (**Supplemental Figure 5e**). Notably, the combination of Avastin or Tivozanib with HEX was well-tolerated, as evidenced by no significant changes in body weight change before and after treatment initiation (**Supplemental Figure 3a,c,e,f**).

### Disruption of TCA cycle anaplerosis, but not enhanced hypoxia, drives the synergy between Avastin and HEX

Hypoxia induction is one of the proximal consequences of angiogenesis inhibition in the intracranial tumors (**Figure 1b-c**). Limited oxygen is known to constrain the amount of ATP produced by oxidative phosphorylation (OxPhos), forcing cancer cells to become increasingly reliant on anaerobic glycolysis to generate ATP^18,19^. Therefore, we reasoned that such metabolic drift to glycolysis could further sensitize *ENO1*-deleted tumors to glycolysis inhibition. *ENO1*-deleted tumor spheres *in vitro* were dramatically more sensitive to POMHEX (a cell-permeable prodrug of HEX) under hypoxic conditions (1% O_2_), as evidenced by a dose-dependent decrease in the TMRE signal, compared to normoxic conditions (21% O_2_) (**Supplementary Figure 6a-d**). Notably, the potency of POMHEX under hypoxia was not limited to *ENO1*-deleted cells as even *ENO1*-intact cells, which are relatively resistant to POMHEX under normoxia, were significantly sensitized to POMHEX under hypoxic conditions (**Supplementary Figure 6a-d**). These findings suggest that hypoxia dramatically enhances the anti-neoplastic efficacy of enolase inhibition by promoting reliance on glycolysis.

Tumor hypoxia is a complex process determined by both demand (consumption) and supply (vascular perfusion) of oxygen to the cells (**Figure 3a**)^36^. Two accurate determinants of tumor hypoxia are: first, oxygen availability (perfusion hypoxia) and second, the rate at which the tumor consumes oxygen in the mitochondria (consumptive hypoxia)^36^. Perfusion hypoxia is dictated by blood perfusion to tumors. Consumptive hypoxia is determined by availability of respiratory substrates such as pyruvate (**Figure 3a**).

**Figure 3:**
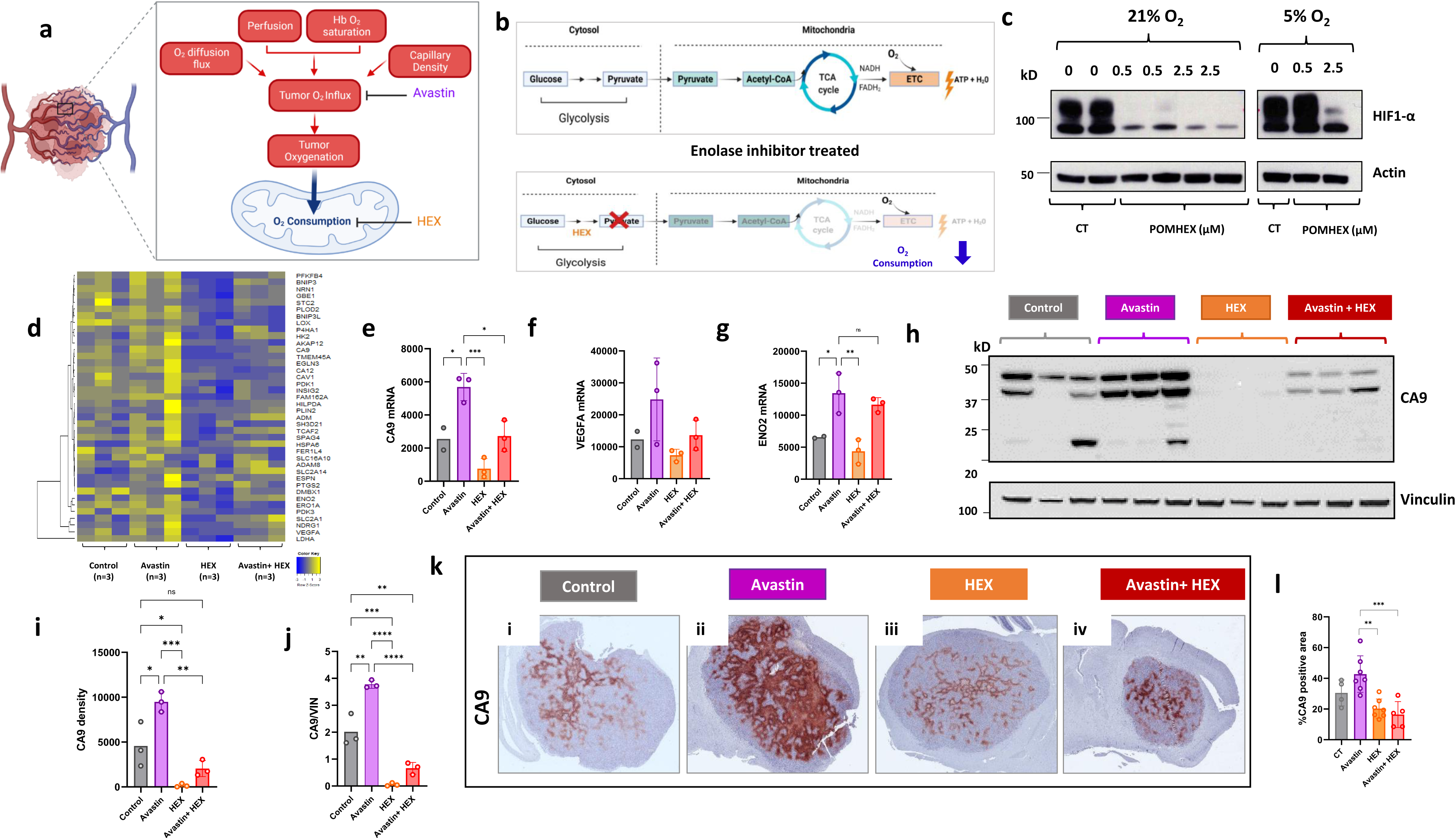
Combination of Avastin and HEX decreases hypoxia in intracranial tumors by limiting oxygen consumption. **a.** Illustrations depicting the roles of vascular perfusion and tumor intrinsic oxygen consuming reactions in overall oxygen tension in the tumors. Hypoxia is a dynamic process and can arise as a result of poor oxygen delivery from the vasculature (perfusion hypoxia) or due to enhanced oxygen consumption by the tumor cells (consumptive hypoxia), or possibly, due to an imbalance in the rates of these two processes. **b.** Schematic demonstrating that mitochondrial oxidative phosphorylation is substrate limited. Inhibition of glycolysis with the POMHEX, a prodrug of Enolase inhibitor HEX, constrains oxidative phosphorylation (oxygen consumption) by blocking formation of pyruvate and decreases consumptive hypoxia. **c**. Western blots showing the effect of glycolysis inhibition by POMHEX on hypoxia (CA9 as a hypoxia marker) in cells *in vitro* in normoxic and hypoxic conditions. **d.** Heatmap representing mRNA transcript levels for hypoxia responsive genes in tumors in each treatment groups. Heatmap shows top 40 hypoxia responsive genes selected from the mRNA profiling studies in the TCGA GBM dataset by correlating CA9 transcript levels (spearman coefficient>0.5) with the whole genome transcripts. **e-g**. Raw mRNA reads of three well-established hypoxia responsive genes, CA9 (**e**), VEGFA (**f**) and ENO2 (**g**) are shown. **h-j**. Immunoblot of intracranial tumors from different treatment groups, showing hypoxia markers CA9 with vinculin as a loading control (**h**). Densitometry analyses of total CA9 signals on each treatment group expressed as total density of signal (**i**) and density of signal relative to the loading control (**j**). **k-l.** Intracranial tumors from mice in control, HEX, Avastin, Avastin and HEX treatment groups, stained with the hypoxia marker CA9 (**k**). Quantification of hypoxic areas (brown) in tumors are plotted for different treatment groups (**l**). Avastin significantly elevates hypoxia in tumors by inhibiting tumor vascularization and impairing perfusion, but HEX treatment reduces hypoxia by inhibiting pyruvate production and suppressing consumptive hypoxia. Avastin and HEX impose a competitive effect on hypoxia, possibly as Avastin elevates perfusion hypoxia while HEX suppresses consumptive hypoxia, leading to a net decrease in intra-tumoral hypoxia. Immunoblot analyses corroborate the results of histopathological analyses, that Avastin and HEX combination significantly decreases hypoxia in intracranial tumors. Data are mean ± SD. Asterik(*) represents statistical significance (p<0.05) achieved by ordinary one-way ANOVA and Tukey’s multiple comparisons test (**e,f,g,h,I,j,l**).

Because HEX inhibits pyruvate production, we first examined the effect of enolase inhibition on hypoxia *in vitro* (**Figure 3b**). Treatment with HEX significantly decreases hypoxia *in vitro* under normoxia and hypoxia, as evidenced by a decrease in the expression of HIF1α and carbonic anhydrase 9 (CA9) and a concomitant increase in mitochondrial OxPhos marker CPT1A (**Figure 3c**, **Supplemental Figure 7a**). One explanation could be that diminished production of glucose-derived pyruvate, a key substrate for mitochondrial oxidative phosphorylation, results in a decrease in consumptive hypoxia^37^ (**Figure 3b**). To further understand the mechanism of synergy of combined angiogenesis and glycolysis inhibition, we performed unbiased transcriptomic analysis on intracranial tumor xenografts treated with HEX and Avastin as mono- or combination therapies. We identified the top 40 genes from the TCGA GBM transcriptomic studies that positively correlated with CA9, a well-known hypoxia marker (Spearman coefficient >0.5; **Figure 3d**). Consistent with our *in vitro* findings, the hypoxia transcriptomic signature is significantly elevated in Avastin-treated intracranial tumors compared to other treatment groups (**Figure 3d-g**). Interestingly, the combination of Avastin and HEX did not significantly alter hypoxia signature (**Figure 3d-g**). This could be explained by a decrease in consumptive hypoxia (mitochondrial oxygen consumption) from enolase inhibition, which negates Avastin-induced perfusion hypoxia (**Figure 3a**). Our *in vitro* observations were corroborated by transcriptomic analysis of hypoxia markers CA9 and VEGFA in intracranial tumor lysates (**Figure 3h-j**) and fixed brain tumor sections (**Figure 3k-l**). Immunoblots on tumor lysates showed an increased CA9 expression in Avastin-treated tumors, a near complete elimination of CA9 expression for HEX-treated tumors, and overall diminished CA9 expression on tumors treated with Avastin and HEX (**Figure 3h-i**). Similarly, immunohistochemistry on fixed intracranial tumors from the Avastin treatment group showed an overall increase in CA9 staining (**Figure 3k-l**). In contrast, tumors from either HEX monotherapy or Avastin + HEX combination therapy produced considerably diminished CA9 expression compared to control or Avastin monotherapy groups (**Figure 3k-l**). Our data reinforce the influence of ‘consumptive’ hypoxia on overall tumor hypoxia and show that ‘perfusion’ hypoxia does not drive the synergy between Avastin and HEX in intracranial tumors in vivo.

Another explanation for the synergy between Avastin and HEX could be the restriction of blood-borne nutrients to tumors caused by angiogenesis inhibition, thereby creating a nutrient-deficient tumor microenvironment (**Supplementary Figure 2c-e**). We previously showed that lower levels of carbon precursors in the medium translate to markedly higher sensitivity to inhibition of glycolysis^37^. Specifically, glycolysis inhibition by (POM)HEX together with low nutrient availability in culture medium, strongly abrogated anabolic reactions crucial for tumor growth^37^. Treatment with Avastin models a nutrient starved tumor microenvironment in vivo (**Figure 1g**). We reasoned that the combination of HEX and Avastin could exacerbate anaplerotic nutrient stress in tumors. As expected, whereas treatment with Avastin or HEX alone modestly depleted TCA cycle intermediates, combined treatment exacerbated this effect (**Figure 5a**). We also found that combined treatment yielded broadly diminished levels of amino acids, including those derived from transamination of TCA cycle intermediates (**Figure 5b**).

Unbiased transcriptomic analysis of the tumors treated with monotherapy of Avastin or HEX, or combination of Avastin and HEX, revealed notable transcriptomic changes. Using DESeq2 analysis, we observed minor changes in differentially up- or down-regulated genes in HEX or Avastin monotherapy groups. In contrast, the combination treatment produced significantly greater changes in differentially expressed genes (DEGs; up: log2Fc ≥; down: log2Fc ≤ −1; padj ≤ 0.05), indicative of a synergistic effect of the drugs on tumor transcriptome (**Figure 4c-f**). For the combination treatment group, genes that overlapped with monotherapy groups also displayed a greater fold change, which supports a synergistic effect of the combination treatment observable at the transcriptomic level (**Figure 4c-f**, **Supplementary Figure 8a-c**). Gene set enrichment analysis (GSEA) further revealed multiple pathways that are enriched in different treatment groups (**Figure 4g-m**). Treatment with HEX monotherapy or combination of Avastin + HEX enriched pathways involved in cellular response to nutrient deficiency—an effect that was exacerbated in combination treatment compared to HEX alone (**Figure 4i, m and Supplementary Figure 9a-b**) which corroborates the metabolomic data (**Figure 4a-b**). GSEA showed statistically significant (q<0.05, NES<-2), negative enrichment of pathways unique to combined treatment, such as those relevant to mitosis, DNA replication, and DNA repair (**Figure 4m**). These observations provide mechanistic support that combined treatment with Avastin and HEX exaggerate anaplerotic nutrient stress and abrogate the anabolic reactions critical for cancer cell proliferation and lead to activation of nutrient stress adaptation response (**Figure 4 and Supplementary Figure 9**).

**Figure 4:**
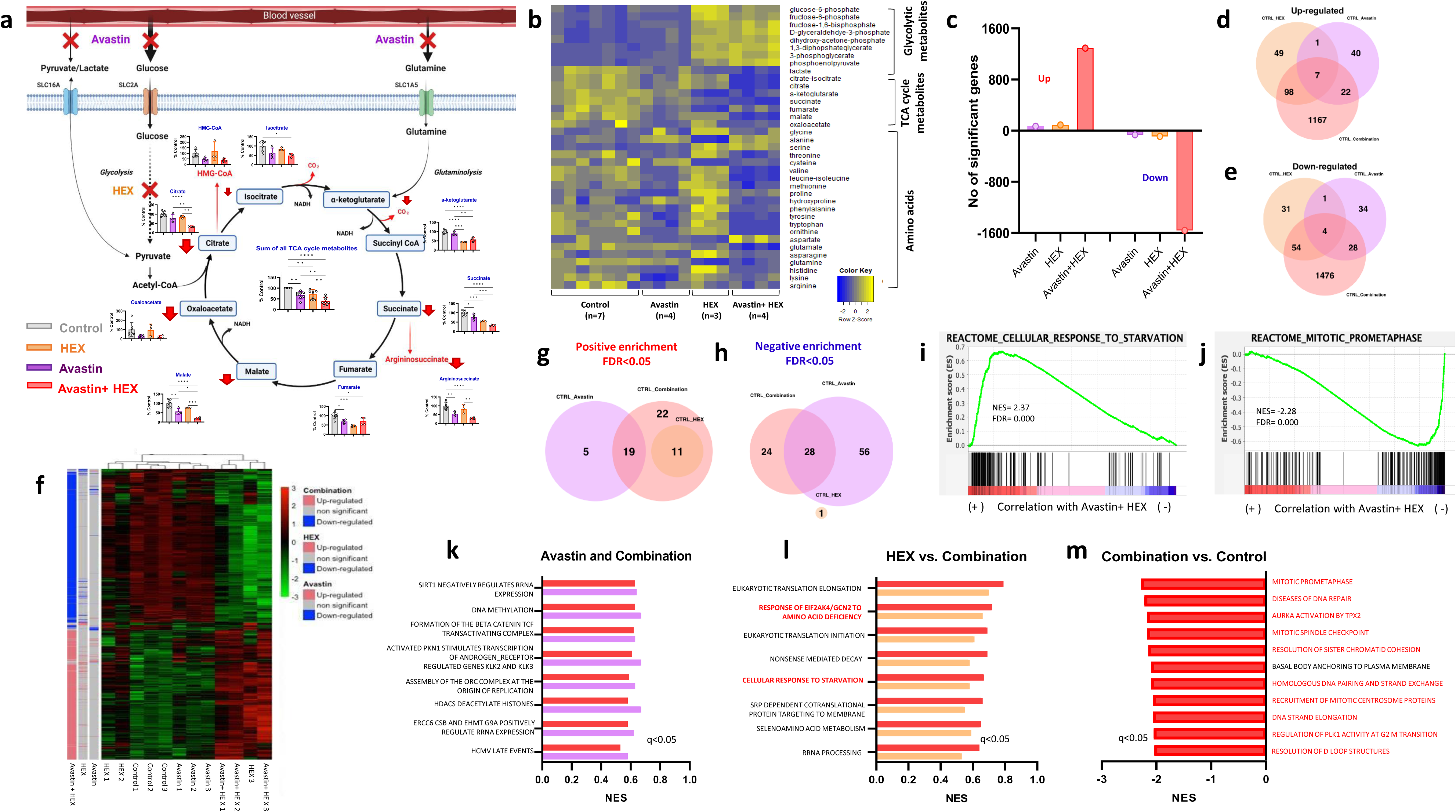
Tumors under combined angiogenesis and enolase inhibition show profound anaplerotic nutrient stress. **a.** TCA cycle map representing metabolites from metabolomic analysis of snap frozen micro-dissected intracranial xenografts of *ENO1* homozygously deleted D423 cells from control, Avastin, HEX or Avastin+HEX groups. Red arrow indicates the degree of decrease in Avastin and HEX treatment relative to control. TCA cycle metabolite depletion achieved by Avastin treatment alone is significantly exaggerated by the addition of the Enolase inhibitor HEX. HEX inhibits anaplerotic pyruvate formation from glucose, and together with Avastin causes greater anaplerotic deficit on the tumors. **b**. Heatmap representing the changes in glycolytic and TCA cycle metabolites and amino acids in the intracranial tumors in control, Avastin, HEX, and Avastin+HEX groups. **c**. Bar-graph showing differentially expressed genes (DEGs) that are statistically significant in different treatment groups (Up: log2Fc ≥ 1 and padj ≤ 0.05; Down: log2Fc ≤ −1 and padj ≤ 0.05). **d-e**. Venn-diagram showing the number of DEGs that are common or unique to each treatment group relative to control. **f**. Heatmap showing statistically significant DEGs in each treatment group. Genes that are up- or down-regulated in each treatment relative to controls are shown. **g,h.** Venn-diagrams representing the gene set enrichment analysis (GSEA) of DEGs relative to control. The number of GSEA reactome pathways that are unique to or overlapping between different treatment groups are shown. **i,j.** GSEA plots showing positive enrichment of genes in the cellular response to starvation pathway and negative enrichment of genes in the mitotic and pro-metaphase geneset. **k,l.** Normalized enrichment score showing the GSEA reactome pathways that are positively enriched (FDR(q)<0.05) in Avastin and Avastin+HEX, and HEX and Avastin+HEX groups. Highlighted in red are pathways that are relevant to cellular nutrient deficiency response. **m**. GSEA reactome pathways that are negatively enriched in Avastin and HEX treatment group relative to control. Highlighted in red are pathways that are relevant in proliferation. Data are mean ± SD. Asterik(*) represents statistical significance (p<0.05) achieved by ordinary one-way ANOVA and Tukey’s multiple comparisons test (**a**).

### Angiogenesis inhibition generates broad metabolic vulnerabilities beyond glycolysis

Given the dramatic synergy between Avastin and HEX in *ENO1*-deleted intracranial tumors, we sought to determine whether other metabolism-targeting therapies would also display similar synergy. Similar to enolase inhibition, inhibition of mitochondrial OxPhos is also marked by depletion of TCA cycle metabolites and most prominently, the amino acid aspartate ^38^. We tested the effects of IACS-010759, an inhibitor of mitochondrial complex 1, on *ENO1*-deleted cells^38^ (**Figure 5**). Previous studies have shown that treatment with IACS-010759 significantly decreases TCA cycle intermediates^38^ (**Figure 5a-f**). Interestingly, toxicity of IACS-010759 in *ENO1*-deleted cells depended on whether cells were cultured in nutrient-rich or nutrient-deplete conditions. Cells grown in nutrient-rich medium treated with IACS-010759 had minimally impacted survival and energy homeostasis, as evidenced by phosphocreatine levels comparable to controls (**Supplemental Figure 10a-d**). However, cells grown in medium with low anaplerotic content were significantly more sensitive to IACS-010759, as evidenced by a significant reduction in phosphocreatine levels—demonstrating exacerbated bioenergetic collapse (**Supplemental Figure 10a-d**). These in vitro observations suggested that in vivo tumors growing in a nutrient-deficient environment induced by angiogenesis inhibition could also be predisposed to sensitization by IACS-010759.

**Figure 5:**
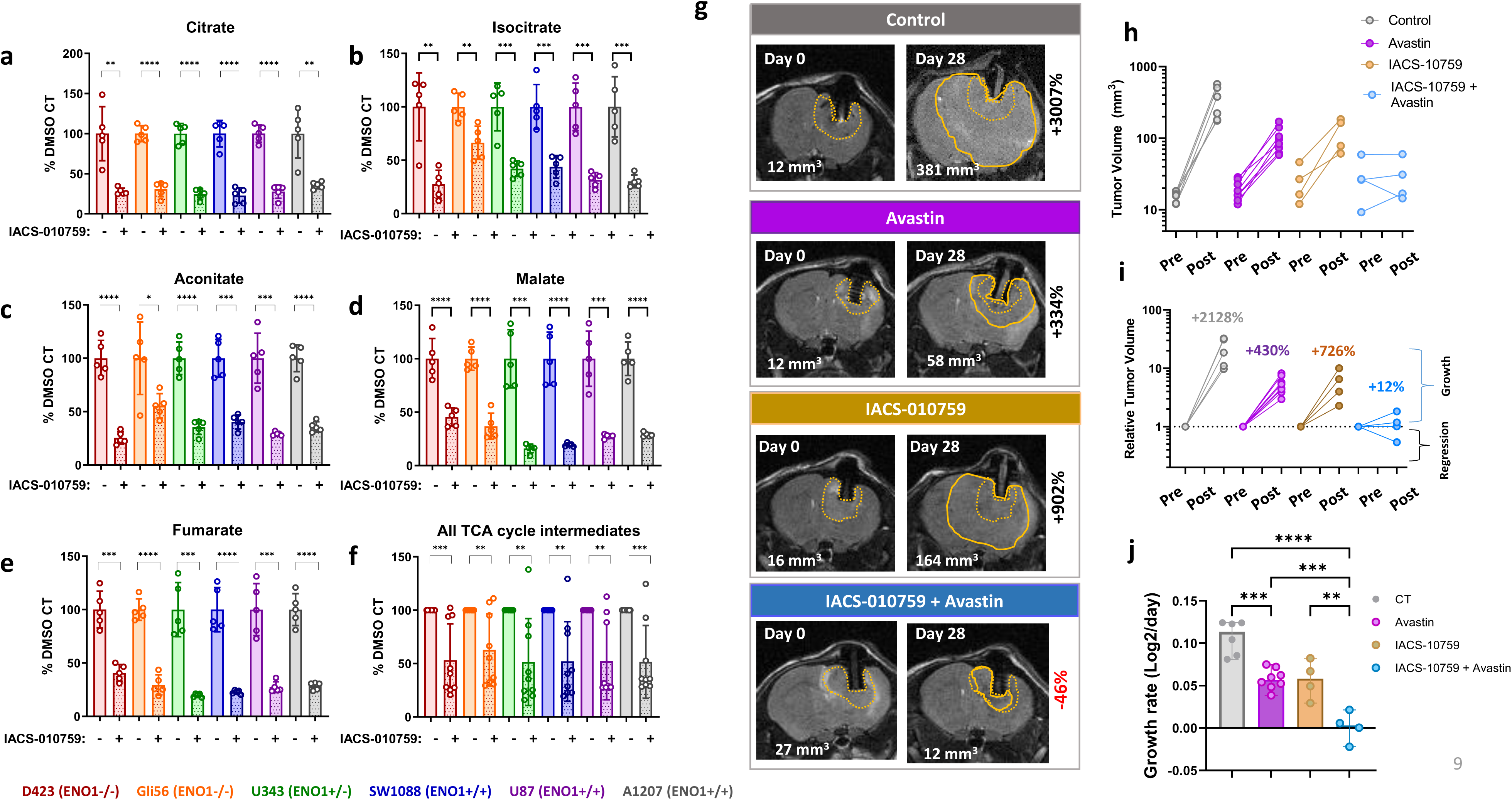
Avastin and the Complex I inhibitor IACS-010759 synergize to abolish grow of *ENO1*-deleted intracranial tumors. **a-f.** Inhibition of oxidative phosphorylation induces anaplerotic stress in cancer cells. Data shown is analyzed from the metabolomic analysis from (Molina et al, Nature Medicine, 2016). Briefly, *ENO1* deleted and *ENO1* intact cancer cells were treated with DMSO or the complex-I inhibitor IACS-010759 and metabolomics was performed to identify different metabolites that are altered in response to mitochondrial complex I inhibition. TCA cycle metabolites are universally diminished by IACS-010759 treatment compared to control in all cell lines irrespective of *ENO1* deletion. **g-j:** Intracranial tumors were generated by implanting *ENO1* deleted glioma cells in Foxn1 ^nu/nu^ nude mice. Tumor development was followed by T2-MRI. Treatment was begun when tumors reached ~20 mm^3^. **g**. T2-MRI images of animals before and after 28 days of treatment with tumor volumes indicated in mm^3^ in the lower part of the image; initial tumor outlines are shown in dotted yellow lines, while tumors after 28 days are shown in solid lines. **h**: Summary of tumor volume changes after 24 days on treatment. Animals were treated continuously with Avastin 2X per week IP, IACS-010759 5 mpk once daily by oral gavage. **i**. Tumor volume plots comparing pre-treatment and 28 days post-treatment raw and relative tumor volumes. **j.** Growth rate of tumor volumes in each treatment group. The effect of IACS-010759 in *ENO1* deleted gliomas is only marginal, but Avastin treatment led to a modest inhibition of tumor growth. Combination of IACS-010759 and Avastin resulted in significant suppression of tumor growth and a complete eradication of tumors in some cases. Data are mean ± SD. Asterik(*) represents statistical significance (p<0.05) achieved by two-tailed unpaired t-test (**a-f**) and ordinary one-way ANOVA and Tukey’s multiple comparisons test (**J**). Note that tumor volume data for *ENO1* deleted tumors in control and Avastin treatment group represented in Figure 5 and Figure 2 are from the same experiment.

*ENO1*-deleted intracranial tumors were treated with either IACS-010759 or Avastin as monotherapies or in combination for 28 days. While treatment with IACS-010759 alone minimally impacted tumor growth in vivo, combined treatment with IACS-010759 and Avastin dramatically delayed tumor growth and produced frank regression of some tumors. (**Figure 5g-j**). These data suggest that angiogenesis inhibition exacerbates metabolic stress and potentiates the anti-neoplastic effect of metabolic inhibitors such as IACS-010759. Additionally, our preliminary experiments also show that that the angiogenesis inhibitor Tivozanib also synergizes with IACS-010759 in xenografts of the NB1 (*PGD*-homozygously deleted) cell line with complete regression observed at doses of IACS-010759 that are ineffective as a monotherapy (**Supplementary Figure 11A-B**).

### Combined inhibition of angiogenesis and metabolism is synergistic against non-glycolysis-compromised orthotopic tumors

We investigated whether inherent compromise of glycolysis was required for Avastin and metabolic inhibitors to exert a synergistic anti-neoplastic effect. Accordingly, we compared the efficacy of Avastin and HEX as single agents or in combination in mice bearing *ENO1*-wildtype (U87) intracranial tumors with the same dosing regimen as we had applied for those with *ENO1*-deleted (D423) tumors (225 mpk BID) for two weeks (**Figure 2**). *ENO1*-deleted tumors showed moderate sensitivity to HEX or Avastin as single agents, and the combined treatment yielded synergistic regression of tumors even with two weeks of treatment (**Figure 6a-d**). However, in *ENO1*-wildtype tumors, with the same treatment regimen and duration, Avastin and HEX as single agents only minimally delayed growth, while combined treatment exerted an enhanced, and possibly additive anti-tumor activity (**Figure 6e-i**). The degree of growth inhibition achieved in *ENO1*-wildtype tumors was dependent on the dose of HEX (**Supplemental Figure 12**). In a separate dose escalation study where HEX was administered 200 mpk three times a day for 8 days, we observed a modest improvement in the efficacy of HEX as a single agent against *ENO1* WT tumors (**Supplemental Figure 12a-c**). Interestingly, the combination of high-dose HEX and Avastin significantly suppressed the growth of *ENO1* WT intracranial tumors with 8 days of treatment (**Supplemental Figure 12a-c**). Histopathological analyses also showed modest decrease in phospho-histone H3 and increase in cleaved caspase-3 signals (**Supplemental Figure 12d-e**).

**Figure 6:**
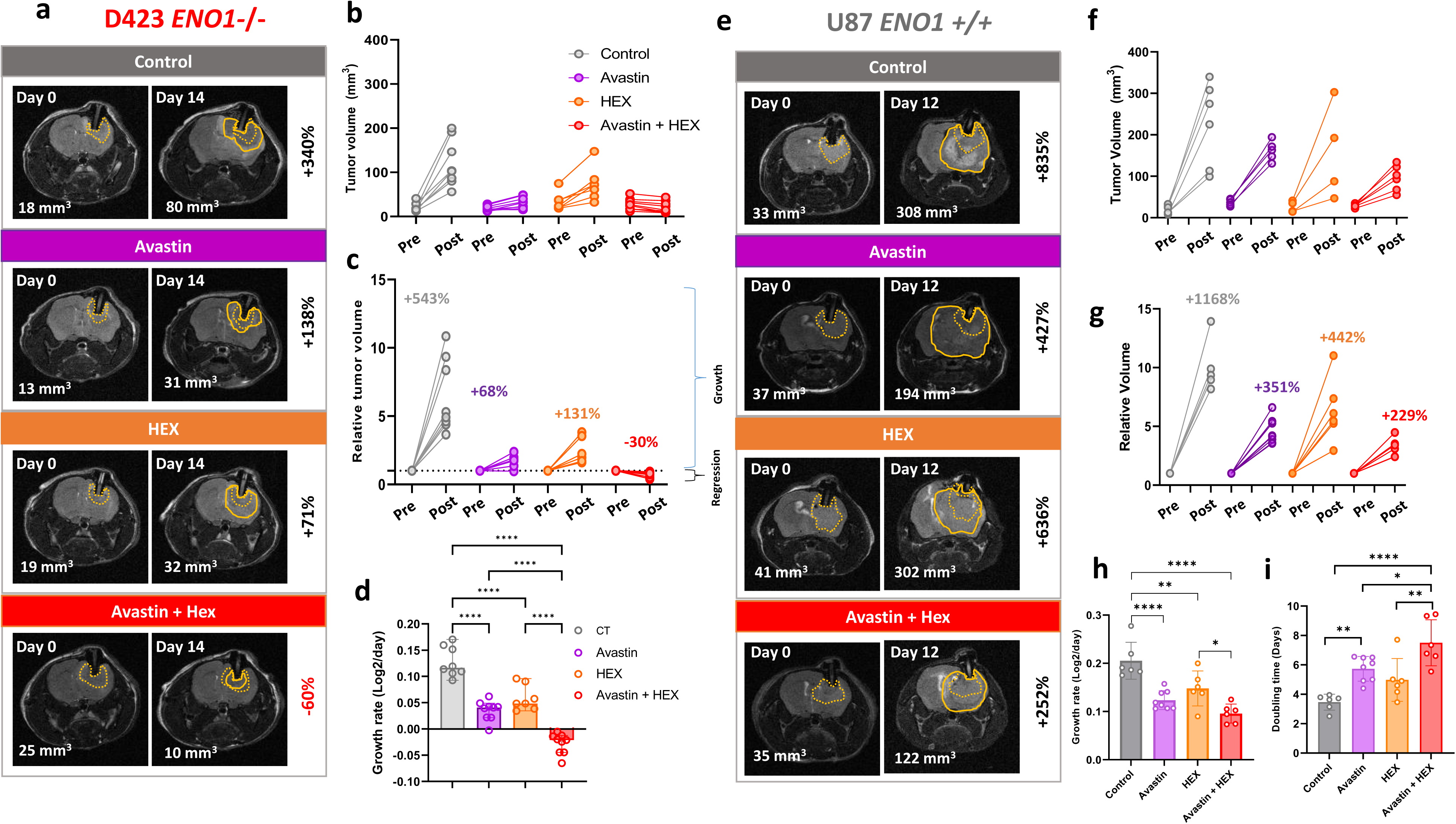
Synergistic anti-tumor effect in *ENO1*-deleted tumors and additive effect in *ENO1*-WT tumors by combination of angiogenesis and enolase inhibitor. Intracranial tumors were generated by implanting *ENO1* deleted D423 and *ENO1* intact U87 glioma cells in immunocompromised nude mice. Tumor development was followed by T2-MRI. T2-weighted MRI images of animal brains with *ENO1* deleted (**a**) or *ENO1* intact (**e**) before and after 14 days (*ENO1* deleted) or 12 days (*ENO1* intact) of treatment with tumor volumes indicated in mm^3^ in the lower left corner of the image; initial tumor outlines are shown in dotted yellow lines, while tumors after 14 or 12 days are shown in solid lines. Animals were separated into four groups—Control (vehicle), Avastin (5 mpk, 2X per week), Enolase inhibitor HEX (225 mpk, 12X per week) or Avastin plus HEX (Avastin, 5mpk 2X per week + 225 mpk SC 12X per week) were administered. **a-d**: Summary of tumor volume changes and growth rates of *ENO1* deleted tumors after 14 days of treatment. **e-i**: Summary of tumor volume changes, growth rates and tumor volume doubling time of *ENO1* intact tumors after 12 days of treatment. HEX or Avastin as single agents suppress tumor growth in *ENO1* deleted tumors; the combination of HEX and Avastin causes a synergistic regression of *ENO1* deleted tumors at the doses administered. In *ENO1* WT tumors, the effect of Avastin and HEX as single agents is marginal, but the combination of the drugs exert an additive effect on tumor growth inhibition. Data are mean ± SD. Asterik (*) represents statistical significance (p<0.05) achieved ordinary one-way ANOVA and Tukey’s multiple comparisons test (**d, h, i**). Note that Day 0 tumor volume data for *ENO1* deleted tumors represented in Figure 2 and Figure 6(a-d) are from the same experiment.

We also tested if the same rationale could be extended to the treatment of *ENO1*-wildtype U87 intracranial tumors with other metabolic inhibitors. Treatment with Avastin sensitized intracranial tumors to IACS-010759, resulting in a significant inhibition of tumor growth, but without frank tumor regression (**Supplemental Figure 13**). Together, these findings indicate that angiogenesis and metabolic inhibitors can work synergistically to inhibit the growth of both glycolysis-compromised tumors and non-glycolysis-compromised tumors, raising the possibility of broad clinical applicability.

### Sensitivity to Avastin is associated with low expression signatures of mitochondrial metabolism and the TCA cycle

Xenografted tumors from different cell lines differ significantly in their response to Avastin and other angiogenesis inhibitors. In the context of this study, we sought to understand the broad molecular features underpinning response to angiogenesis inhibitors—specifically, whether anaplerotic nutrient stress or diminished mitochondrial metabolism is a determinant of sensitivity to Avastin in human tumors. To that end, we interrogated a set of PDX models from CrownBio that had corresponding transcriptomics and data on anti-tumor response to Avastin (**Supplemental Figure 14**); transcriptomic analyses were performed on tumors prior to Avastin treatment initiation. We searched for specific transcriptomic characteristics that correlated with anti-tumor responsiveness across a range of independent PDXs. Our analysis revealed a strong negative correlation between Avastin sensitivity and mitochondrial metabolic gene signature in tumors at baseline (**Supplemental Figure 14-c**). More specifically, tumors possessing elevated gene signature for mitochondrial metabolic processes, such as mitochondrial complex I biogenesis, ATP biosynthesis, TCA cycle and mitochondrial respiratory chain reactions, better tolerated Avastin treatment relative to PDXs with lower mitochondrial metabolic gene signature (**Supplemental Figure 14b-c**). We also found that tumors with enrichment in gene signatures that bolster response to nutrient deficiency at baseline, were less sensitive to Avastin reinforcing the notion that nutrient stress adaptation may predict Avastin sensitivity (**Supplemental Figure 14b**). These PDX findings complement our key observation that anaplerotic nutrient stress achieved by treatment with either glycolysis inhibitor HEX or mitochondrial OxPhos inhibitor IACS-010759 sensitizes tumors to angiogenesis inhibition.

## DISCUSSION

Here, we sought to uncover potential angiogenesis-inhibition induced metabolic vulnerabilities. Comprehensive metabolomic and transcriptomic analysis of intracranial tumor xenografts revealed that anti-angiogenic therapy generates an anaplerotic nutrient deficit and sensitizes xenografted tumors to inhibitors of key energy metabolism pathways, such as glycolysis and oxidative phosphorylation. Combined treatment of the angiogenesis inhibitor Avastin and the glycolysis inhibitor HEX yielded synergistic anti-tumor effects accompanied by dramatically exacerbating metabolomic and transcriptomic signatures of anaplerotic nutrient stress in intracranial tumors.

The blood-brain-barrier is poorly permeable to small hydrophilic molecules such as HEX and an initial explanation for its efficacy as a monotherapy would have posited permeation through the tumor-associated leaky vasculature. The dramatic synergy between Avastin and HEX, despite Avastin’s effect on normalizing (sealing) the BBB, indicates that HEX must be reaching the malignant glioma cells through another mechanism. Low molecular weight phosphonate drugs are known to reach the brain through fenestrations in the blood-CSF barrier^32,39^. Unlike the BBB, capillaries in the blood-CSF barrier have fenestrations, which allow low molecular weight, hydrophilic drugs to permeate into the brain; one prominent example of such is the antibiotic fosfomycin, which is used clinically to target brain abscesses (**Supplementary Figure 4b**)^39^. Given the high physiochemical similarity – it is most likely that HEX reaches brain and brain tumor through permeation via the blood-CSF barrier and the Avastin-induced re-sealing of the BBB is inconsequential for HEX entry (**Supplementary Figure 4b**). Furthermore it is worth noting that the combination of HEX and angiogenesis inhibitors may yield superior results in humans given that HEX has a an unusually short half-life in mice; with the drug being undetectable in plasma in as little as 2 hours after injection^33^. This is in contrast to rat, dog and non-human primate where the half-life of HEX is ~10-fold longer^33^.

We also showed that Avastin treatment sensitizes *ENO1* deleted tumors to inhibition of mitochondrial oxidative phosphorylation by IACS-010759. Previous studies reported that homozygous deletion of *PGD* causes even stronger OxPhos inhibitor vulnerability^38,40^. Accordingly, we showed that angiogenesis inhibitor Tivozanib also synergizes with IACS-010759 in xenografts of the NB1 (*PGD*-homozygously deleted) cell line with complete regression observed at doses of IACS-010759 that are ineffective as monotherapy. Importantly, studies have suggested a therapeutic benefit in combining Avastin with the mitochondrial complex I inhibitor, metformin for the treatment of recurrent type I endometrial cancer, ovarian cancer, and non-small cell lung cancer^41–43^. Our findings are consistent with this line of reasoning and provide a mechanistic basis for the efficacy of the combination treatment modality. Finally, combined treatment with Avastin and HEX/IACS-010759 also showed therapeutic benefit against glycolysis intact wild-type tumors, raising the possibility of therapeutic relevance across a broad range of malignancies. With respect to clinical application, it is worth noting that we can achieve significant regression of tumors at doses of IACS-010759 which are far lower than those employed in published literature^38^. This is important since inhibition of mitochondrial OxPhos yielded grade 3 peripheral neuropathy in recent phase 1 trials: 11/23 patients in the solid tumor trial and 4/11 patients in the relapsed/refractory acute myeloid leukemia trial (NCT02882321, NCT03291938)^44^. A dose-reduction of IACS-010759 enabled by combination with angiogenesis inhibitors may allow resuscitation of this drug given its very sharp dose-limited toxicity^44^.

To definitively correlate our pre-clinical findings with human clinical data, it would be highly desirable to perform a thorough metabolomic profiling of primary human tumors subjected to angiogenesis inhibitors. However, due to hemorrhagic complications and challenges associated with wound healing, anti-angiogenic therapies preclude surgical resection of tumors (treatment discontinuation required for at least 28 days before surgery)^1^, which restricts a systematic assessment of metabolic alterations caused by angiogenesis inhibition in human tumors. This may also explain why, despite the widespread use of angiogenesis inhibitors in the clinic, limited investigations have been performed to understand how these drugs affect the transcriptomic or metabolic landscape of human tumors. We circumvented this key limitation by employing human cell line-derived xenograft (CDX) as well as patient-derived xenografts (PDX). Using different - omic profiling approaches, we identified crucial, exploitable vulnerabilities associated with angiogenesis inhibition. Overall, our data strongly suggests that anaplerotic nutrient stress incurred by tumors due to angiogenesis inhibition enhances sensitivity to therapies targeting tumor metabolism.

In conclusion, our findings reinforce clinical evidence showing that Avastin, despite significantly reducing blood flow to the tumors, only temporarily halts tumor growth. However, essential metabolic adaptations that arise in the context of angiogenesis inhibition can be leveraged to sensitize tumors to different metabolism-targeting therapies. Our preclinical data strongly invite a systematic investigation of combinations of angiogenesis inhibitors with therapies targeting cancer-specific metabolic vulnerabilities beyond HEX and IACS-010759. For instance, therapeutic agents that target pathways converging on anaplerosis, such as glutaminolysis inhibitors (glutaminase inhibitor CB-839, the ASCT2 inhibitor V-9302^45^), or other anabolic and catabolic pathways involving the metabolism of asparagine, (L-arsparaginase^46^) methionine (MAT2A inhibitor AG-270, PRMT5 inhibitor GSK3326595^47^ and arginine (ADI-PEG20^48,49^) (**Figure 7**). An additional mechanism worth investigating may be interactions with metabolic therapies involving caloric restriction or specific macronutrient intake, such as ketogenic diets which restrict carbohydrates, which may further enhance the therapeutic efficacy of anti-angiogenic therapies. Our paper thus demonstrates the mechanistic basis for combining anti-metabolic and anti-angiogenesis therapies to potentiate anti-tumor efficacy and potentially improve overall patient survival.

**Figure 7:**
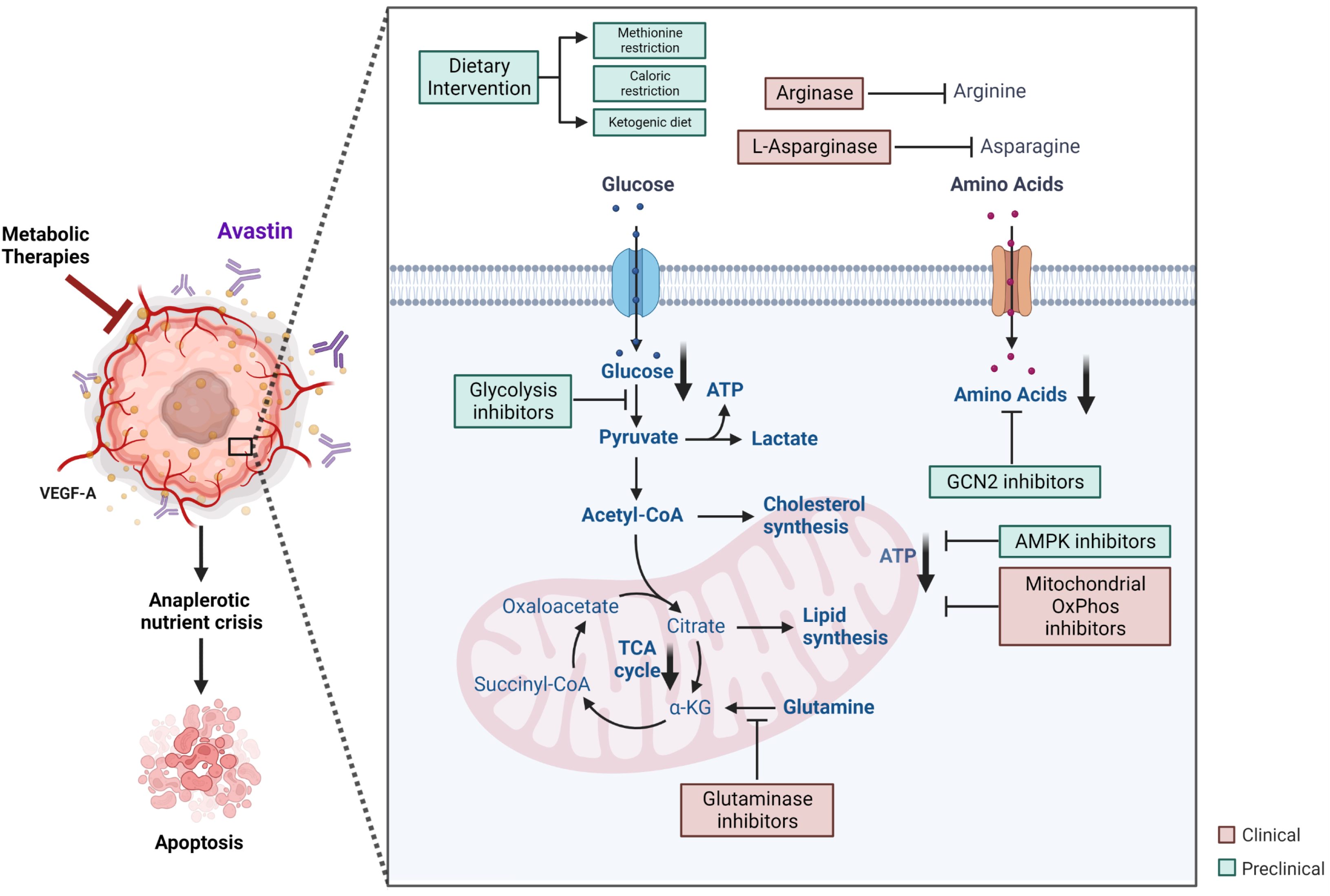
Potential of angiogenesis inhibitors as adjuvant/combination treatment with metabolic therapies in cancer. Illustration highlighting synergistic anti-tumor therapeutic potential of combining angiogenesis inhibitors with existing and emerging metabolic therapies that result in anaplerotic nutrient stress in cancer cells. Angiogenesis inhibitors such as Avastin impede the delivery of blood-borne nutrients such as glucose and amino acids to cancer, resulting in anaplerotic deficit, and expose a multitude of metabolic vulnerabilities in cancer cells. Such metabolic liabilities may be exploited further through inhibition of glycolysis (POMHEX/HEX), as well as amino acids availability/breakdown (L-asparginase, Arginase, CB-839), impairment of metabolic stress adaptation (GCN2 inhibitors, AMPK inhibitors, autophagy inhibitors), and integration of dietary interventions such as methionine restricted diet, caloric restriction, and ketogenic diet. Together, the combination of angiogenesis inhibition and metabolic therapies may accentuate anaplerotic nutrient crisis and yield a synergistic inhibition of cancer cell growth and proliferation.

## METHODS

### Cell Lines

The H423/D423-MG (CVCL_1160, glioblastoma) cell line, which has a 1p36 homozygous deletion spanning *ENO1*, was generously provided by D. Bigner^50^. D423 *ENO1*, an isogenic cell line with ectopic constitutive expression of *ENO1*, was generated by our lab as described previously. The *ENO1*-WT cell lines LN319 (CVCL_3958, a **sub-clone** of LN-992) and U87 (CVCL_0022) were obtained from the Department of Genomic Medicine/IACS Cell Bank at MD Anderson.

All cells were used below passage 25 and maintained in DMEM (pH 7.4) supplemented with 4.5 g/L glucose, 110 mg/L pyruvate, and 584 mg/L glutamine (Cellgro/Corning #10-013-CV) at 37°C in a 5% CO2 atmosphere unless indicated otherwise. DMEM was supplemented to achieve a composition of 10% fetal bovine serum (Gibco/Life Technologies #16140-071), 1% PenStrep (Gibco/Life Technologies #15140-122), and 0.1% amphotericin B (Gibco/Life Technologies #15290-018). Cell lines were regularly checked for mycoplasma contamination with the MycoAlert PLUS detection kit (Lonza) and authenticated by STR fingerprinting with the Promega PowerPlex 16 System. STR fingerprinting was conducted by personnel in MD Anderson’s Characterized Cell Line Core (CLCC). STR profiles were compared to both the CLCC database and external cell databases (DSMZ/ATCC/RIKEN/JCRB).

### Intracranial orthotopic tumor cell implantation

All experiments involving mice were performed at MD Anderson under a protocol approved by MD Anderson’s Institutional Animal Care and Use Committee (IACUC).

Intracranial glioma tumors were generated by injecting 200,000 cells into the brains of 4- to 6-month-old immunocompromised female nude Foxn1nu/nu mice, which were bred at MD Anderson’s Experimental Radiation Oncology Breeding Core. Prior to intracranial tumor cell injection, a bolt (a plastic screw with a hole in its center) was drilled into the skull of each animal. The animals were allowed to recover for 2 weeks, during which time they were monitored for signs of morbidity. Two weeks after bolt implantation, the cells were injected through the bolt using a Hamilton syringe. Animals exhibiting any severe neurological morbidities were after tumor implantations were euthanized. Intracranial bolting and injections were performed by personnel in MD Anderson’s Intracranial Injection Fee-for-Service Core^51^. All procedures were performed in accordance with the regulations of MD Anderson’s IACUC.

### Tumor volume measurement in vivo

Mice bearing intracranial tumors underwent T2-weighted MRI weekly on a 7T Biospec USR707/30 scanner (Bruker Biospin MRI, Billerica, MA) in MD Anderson’s Small Animal Imaging Facility. Prior to imaging, the animals were briefly anesthetized with isoflurane. Throughout the imaging procedure, the animals’ body temperatures were maintained with a heating blanket; their bodies and heads were restrained with a stereotactic holder; and their heart and breathing rates were monitored.

For T2-weighted MRI, a low-resolution axial scan was first taken to calibrate the scanner position. Then, two high-resolution axial scans and one high-resolution coronal scan were taken. For the axial scans, the slice thickness was 0.500 mm, and the increment between each slice was 1.000 mm. To obtain better tumor coverage, we offset the two axial scans by 0.500 mm, which was equal to the difference between the slice increment and slice thickness. The coronal scans had a slice thickness of 0.750 mm and a slice increment of 1.000 mm. The total number of slices for each scan was based on the size of the tumor. The MRI scans were analyzed with the open-source image-processing software program 3D Slicer (v4.10, http://www.slicer.org)^52^.

MRI series were independently reviewed, and tumor volumes determined in 3D Slicer, by three lab members. The draw tool in the editor module of 3D Slicer was used to manually delineate tumor tissues slice-by-slice. Enhanced contrast due to edema, as well as hollow areas of tumor caused by bolting, were excluded. The tumor volume of each scan was calculated automatically by using the Label Statistics module, which calculates the tumor volume for a scan by converting the selected pixel area on each slice to squared centimeters, multiplying that value by the slice increment, and then summing up the slice volumes. The tumor volume for each mouse was calculated as the average of the three scans (2 sets of axial scans and one set of coronal scans). Finally, all three independent measurements were averaged.

### Polar metabolite profiling

The mice were euthanized following the standard IACUC protocol. The animals were then dissected to extract subcutaneous tumors. The weights of the tumors were recorded, and the tumors were cut into two pieces, one of which was snap-frozen in liquid nitrogen for metabolomic analysis. The tumor pieces snap-frozen in liquid nitrogen were cut into small pieces (~50 mg) and placed into pre-cooled microcentrifuge tubes (Fisher Scientific, cat. 02-681-291) containing steel beads (Qiagen). To these tubes, we added 1 mL of 80% methanol pre-cooled to −80 °C. Tumor tissue was bead-mill homogenized with the Qiagen TissueLyser by shaking tubes at 28 Hz for multiple rounds of 45 s each. To the tubes of the homogenized tumor lysates, we added 80% methanol to create a final weight-adjusted volume of 25 mg/ml. After incubation for 15 min on dry ice, tumor lysates were homogenized once more using a vortex mixer for 1 min. The samples were then centrifuged for 5 min at 14,000 x g at 4 °C. Cell debris and non-polar metabolites precipitated and were separated from the polar metabolites, which collected in the supernatant. The polar metabolites were transferred into chilled 1.5-ml Eppendorf tubes and dried in a SpeedVac (Thermo Fisher). The dried and concentrated polar metabolites were submitted to John Asara’s Polar Metabolite Profiling Platform at Beth Israel Deaconess Medical Center. There, the samples were subjected to tandem mass spectrometry via selected reaction monitoring with polarity switching for 300 total polar metabolite targets using a 5500 QTRAP hybrid triple quadrupole mass spectrometer (SCIEX). The mass spectrometer was coupled to a high-performance liquid chromatography system (Shimadzu) with an amide hydrophilic interaction liquid chromatography column (Waters) run at pH=9.0 at 400 mL/min. Q3 peak areas were integrated using MultiQuant 2.1 software^32^.

### Immunohistochemistry

Tumor-bearing mice were euthanized at the end of the experiment or earlier if they exhibited neurological symptoms, and their brains were harvested and then fixed in 10% paraformaldehyde. Mouse brains were dissected and submitted to MD Anderson’s Veterinary Pathology Core for dehydration, paraffin embedding, and tissue sectioning. Tissue sections were dried overnight at 60 °C and then deparaffinized in xylene. The deparaffinized sections were rehydrated by a series of washes in aqueous solutions of decreasing ethanol concentration. Antigens were retrieved by boiling the sections in citrate buffer (1:100 Vector Antigen Unmasking Solution [Citrate-Based] H-3300 250 mL) for 10 min and then cooling the sections for 30 min. Tissue sections were blocked with 2% goat serum (Vector S-1000 Normal Goat Serum; 20 mL) in PBS (Quality Biological PBS [10X], pH 7.4; 1000 mL) for 1 h. The sections were then incubated with monoclonal anti-cleaved caspase 3 rabbit (cleaved caspase-3 (Asp175) (5A1E) rabbit mAb; CST# 9664T, Cell Signaling Technology) or anti-phospho S10 histone H3 (rabbit anti-phospho histone H3 [S10] IHC antibody, affinity purified; Bethyl Laboratories, IHC-00061 or rabbit anti-carbonic anhydrase 9 antibody (CST#5649)) diluted to 1:1000 with 2% goat serum in PBS overnight at 4°C. The next day, the sections were washed three times with PBS in a shaker. After washing with PBS, the sections were incubated with 1X goat anti-rabbit IgG poly HRP secondary antibody (Invitrogen by Thermo Scientific) for 30 min and then washed in PBS and Tween 20 (Fisher BioReagents BP337-500). The sections were stained with either ImpactNOVAred (Vector Labs SK-4805) or EnzMet^tm^ (Nanoprobes #6001-30 mL) and then counterstained with hematoxylin or with hematoxylin and eosin, respectively. The stained sections were mounted using Denville Ultra Microscope Cover Glass (#M1100-02) and Thermo Scientific Cytoseal 60 and dried overnight at room temperature.

### RNA sequencing, and data analysis from treated xenografted tumors

Tumors were extracted from frozen brain sections from mice treated with vehicle, Avastin, HEX or Avastin + HEX for 14 days. Briefly, whole brain was dissected out from the mice following euthanasia and immediately snap frozen in liquid nitrogen. Xenografted tumors from frozen whole brain was carefully cut out as 2 mm wide cross-sectional slices. Axial MRI scans were used as guide to cut xenografted tumor from the whole brain. For RNA extraction from the tumors, approximately 1ml of Trizol reagent was added to 5 mg of frozen xenografted tumors and homogenized using a homogenizer (POLYTRON® PT 1200E, Kinematica #9112212) 200 µl chloroform was added to the homogenized tumors in Trizol, and after a vigorous shaking of the tubes, phase separation was performed by centrifugation. Aqueous phase solution was precipitated with isopropanol, and RNA purification was performed using the Qiagen RNA extraction kit (steps 4-8). Approximately, 200 ng of RNA was submitted to BGI to perform 100 bp paired end for 30 million transcript reads. The FASTQ files obtained from BGI were aligned with hg19 to generate sam files using the HISAT/StringTie/Ballgown modules^53^. Differential gene expression between the control and experimental groups were determined by using the edgeR package v3.34.0, To identify significant differentially expressed genes, we used the following criteria: log_2_ fold change (log_2_Fc) ≥ 1 (for the Up-regulated gene) or log_2_Fc ≤ −1 (for the down-regulated gene), and an adjusted p-value (padj) ≤ 0.05. Reactome pathway enrichment was determined using the gene set enrichment analysis (GSEA)^54^ with default parameters and Reactome subset (c2.cp.reactome.v2022.1.Hs.symbols.gmt from Human MSigDB Collections)^55^. Heatmaps were generated using the ggplots or pheatmap package in R v4.0.3.

### Patient-derived xenograft (PDX) models (Crown Bio datasets)

PDX models were developed and established in immunodeficient mice at Crown Bioscience as previously described^56^. Briefly, cryopreserved or fresh tumor tissues were cut into small pieces (~2-3 mm in diameter), and subcutaneously transplanted in the right flank of NOD/SCID or BALB/c nude mice. Tumor growth was checked twice a week using a caliper. Pharmacological dosing started when tumor size reached 100-300 mm^3^. Tumor-bearing mice were euthanized when tumor size reached 3000 mm^3^. All procedures were performed in pathogen-free animal facility at Crown Bioscience under the approved protocols by the Institutional Animal Care and Use Committee.

### RNA-seq

Library construction for RNA-seq was performed using MGIEasy RNA Library Prep Set (MGI, catalog no. 1000006383) following the manufacturer’s instructions. Briefly, poly-A mRNA was captured from total RNA using Oligo-dT-attached magnetic beads (Agencourt, catalog no. A63987) and fragmented. cDNA was synthesized and purified, followed by A-tailing and ligation of adapters. DNA fragments with adapters were selected and amplified by PCR. The quality of the final library was checked by Qubit and Agilent 2100 Bioanalyzer. Paired-end sequencing with a read length of 150bp was performed following the manufacturer’s instructions (MGI, catalog no. 1000012555).

### Alignment and quantification of transcripts

The quality of raw data was checked by FastQC (version 0.11.9). Adapter and low-quality sequences were trimmed by Trimmomatic software (version 0.40)^57^. Sequencing reads were mapped to human (hg19) reference genomes using STAR (version 2.7.10a)^58^, based on which alignment yielded fewer mismatches, they were subsequently sorted into human or mouse groups, representing cancer or stromal transcriptomes, respectively. Ambiguous reads were discarded. Transcript-level read counts were quantified using kallisto (version v0.46.1)^59^; and subsequently summarized to gene-level count and abundance estimates. The count estimates were used as inputs in differential expression analysis (see **Differential expression analysis on CDX dataset**). The gene-level transcript abundance values (in transcripts per million, or TPM) were used as inputs in biomarker discovery (see **Biomarker discovery in PDX dataset**).

### Differential expression analysis on CDX dataset

Differential expression analysis was performed using DESeq2 (version 1.34.0)^60^ cancer (human) and stroma (mouse) genes. The following generalized linear model was fit to explain the expression level of each gene with respect to treatment groups, accounting for possible interaction between Avastin and HEX treatments:

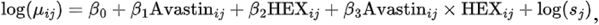

where read count for gene *i*, sample *j* is assumed to follow a negative binomial distribution with fitted mean *μ_ij_* and sample-specific size factor *s_j_*. Regression coefficients *β_1_* and *β_2_* measure the effects of Avastin and HEX treatments on gene expression, and *β_3_* measures how the effect of Avastin treatment changes when HEX treatment is also present.

### Biomarker discovery in PDX dataset

The Crown Bioscience PDX dataset consist of tumor volume records in 21 PDX models under Avastin and negative control treatments, as well as the cancer transcriptomic profiles of these PDX models. The 21 PDX models used in this study comprise following cancer types: breast (n = 1), cervix (n = 4), cholangiocarcinoma (n = 2), colon (n = 7), kidney (n = 1), liver (n = 2), lung (n = 2), and ovary (n = 2). The following linear mixed model was used to explain tumor volume at day *t* for mouse *i* in PDX model *j*:

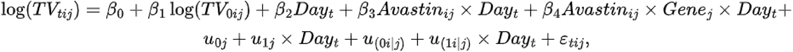

where *t*, *i*, *j* indicates time point, mouse, and PDX, respectively. ‘Gene’ is a covariate for gene-level transcript abundance values in log scale. For a detailed description of the fixed and random effect terms, please refer to Guo *et al*^56^. Using Benjamini-Hochberg adjusted p-value < 0.05 as cut-off, we obtained a list of genes whose expression levels significantly impact the efficacy of Avastin treatment (*β_4_* ≠ 0). This gene list was used as input in gene set enrichment analysis.

### Gene set enrichment analysis

Gene set enrichment analysis (GSEA)^54^ performed using clusterProfiler (version 4.2.2)^61^.The input gene lists were ranked based on Q = sign(*β*)·log(*p*), where *β* and *p* are the regression coefficient and its associated p-value obtained from the differential expression and biomarker analyses described in previous sections. Gene Ontology and REACTOME gene sets were retrieved using msigdbr (version 7.5.1).

### Western blots

#### Cells

Cells were grown in 6 well plates and treated with POMHEX for 48 hours in hypoxic (5% O_2_) and normoxic conditions (21%). Whole cell lysates were harvested by washing the cells twice with ice cold phosphate-buffered saline (PBS). RIPA buffer (ice-cold) was supplemented with protease (cOmplete™ mini, Roche#11836153001) and phosphatase inhibitors (PhosSTOP, Roche, #5892970001), added and the samples were then sonicated.

#### Tumors

Frozen intracranial tumors were cut into smaller pieces and approximately 20 mg of tumors was used to prepare lysates. RIPA buffer supplemented with protease and phosphatase inhibitors was added to the tumor samples and thoroughly homogenized. The tumor homogenates were further sonicated and centrifuged.

The protein concentrations in cell/tumor lysates were determined using the BCA assay (ThermoFisher, #23227). After being separated by Nu-PAGE SDS-PAGE (4-12% gradient) (TransBlot turbo), proteins were deposited onto nitrocellulose membranes using the semi-dry technique. The successful transfer of proteins onto the membrane was confirmed with Ponceau S staining. To block the non-specific membrane sites, 5% non-fat dried milk in tris-buffered saline (TBS) with 0.1% Tween 20 (TBST) was used as the blocking agent. The membranes were treated with primary antibodies overnight at 40C with moderate rocking. The membranes were TBST-washed three times for five minutes on the second day. After that, membranes were gently rocked for an hour while incubating in HRP-tagged secondary antibody (1:5000). The antibodies used in our investigations were: (HIF1-α (CST #14179), β-actin (CST#3806), CA9 (CST#5649), Vinculin (CST#13901), Atg5 (CST#9980), LC3B(CST#3868), pAkt (CST#9271), Akt (CST#9272), CPT1A (CST#12252), anti-rabbit HRP linked antibody (CST#7074), anti-mouse HRP linked antibody (CST#70776)).

### Statistical analysis

Statistical analyses reported in this study were performed using either Microsoft Excel or Graph Pad Prism 8. Unpaired Student’s test and 1- or 2-way ANOVA were used where appropriate. Tukey’s post hoc analysis was used to determine statistical significance following ANOVA. P<0.05 was used as a threshold to determine statistical significance.

## Illustration

All illustrations were made using BioRender.

## Author’s contributions

SK and FLM conceived the study and designed the experiments. SK and YL performed all intracranial animal experiments, including drug treatments and MRI imaging, with help from KA and SC. SK, YH, YC, EB, AP, and TN calculated tumor volumes from MRI scans. SK and FLM analyzed the data from all *in vivo* experiments. JA, AP, and TN performed immunohistochemistry and EC scanned the histology slides. YH performed *in vivo* metabolomics experiments and JA performed mass-spectrometry analysis. SK, YC, WH, FB, YS, and SG analyzed the transcriptomic data. MNU synthesized HEX for animal treatments. DG assisted with the logistics for animal experiments, data analysis and figure preparations. SM provided financial support, and RAD provided key insights on the study and supervised experiments. SK wrote the manuscripts with assistance from FLM and VCY. All authors approved the final manuscript.

## Acknowledgements

We thank Charles Kingsley and the team from MD Anderson SAIF for their assistance with MRI scans of intracranial tumors. We thank the Editing Services team at Research Medical Library at the University of Texas MD Anderson Cancer Center for the editorial support on our manuscript. We thank Michael Hazel from Crown Bioscience, Dr. Seth Gammon, Dr. Brian Engel from MD Anderson Cancer Center for helpful discussions pertinent to our investigation.

---

**Supplementary Figure 1.**
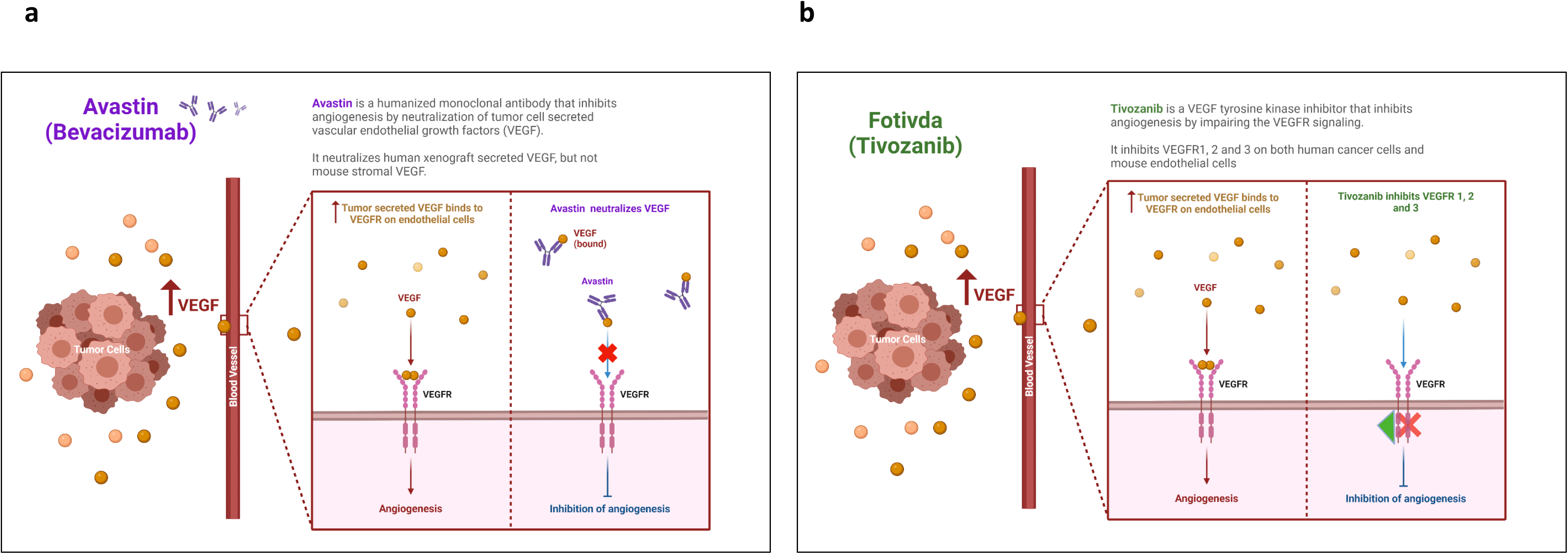
Angiogenesis inhibitors impair neo-vascularization by inhibition of the VEGFR signaling cascade. **a.** Bevacizumab (Avastin) is a VEGF-A neutralizing humanized monoclonal antibody that neutralizes the VEGFA secreted by the tumor cells and impairs VEGFR signaling (Ferrara et al. 2004 *Nat Rev Drug Discov.*). Avastin is clinically approved as a frontline therapy for a broad range of malignancies that include glioblastoma multiforme, metastatic colorectal cancer, renal cell carcinoma, ovarian adenocarcinoma among many others. **b.** Tivozanib (Fotivda) is a specific inhibitor of VEGFR 1, 2 and 3 and it abrogates the VEGF signaling pathways through the inhibition of the tyrosine kinase activity of the VEGFRs (Nakamura et al. 2006 *Cancer Res.*).

**Supplementary Figure 2.**
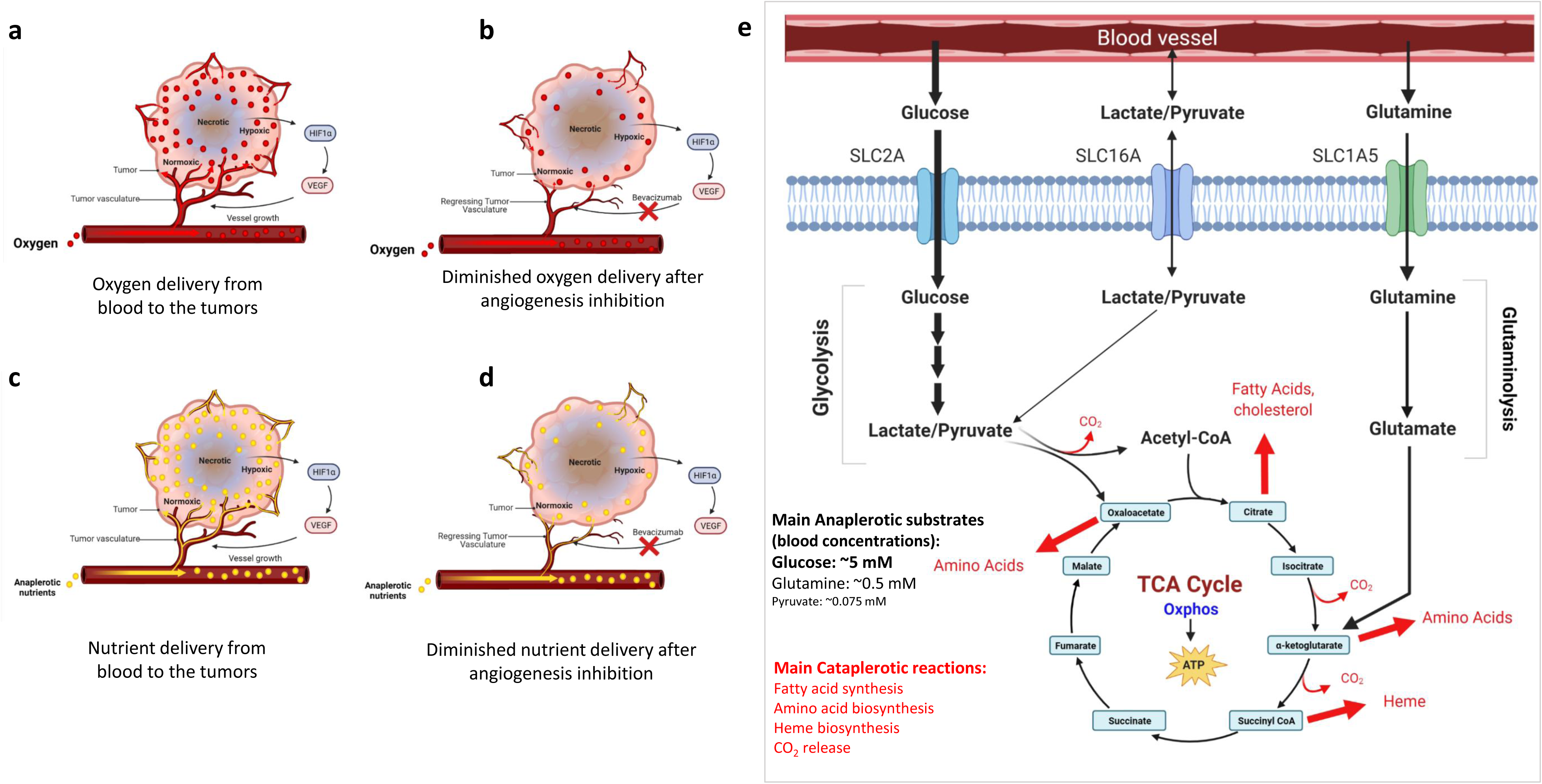
Angiogenesis inhibition impairs neovascularization in tumors, restricts tumor perfusion and delivery of oxygen and blood borne anaplerotic nutrients/Oxphos substrates. **(a-d)** Schematic depicting the impact of tumor perfusion on oxygen and nutrient delivery to the tumors following angiogenesis inhibition. Angiogenesis ensures a continuous delivery of oxygen(**a-b**) and import of nutrients(**c-d**) to the tumor cells enabling metabolic processes that generate the biosynthetic and bioenergetic intermediates to support tumor survival and proliferation. Tumors employ multiple signaling pathways, most notably, HIF1α pathway, once they sense a hypoxic environment. One of the direct consequence of HIF1α activation is the secretion of vascular endothelial growth factors (VEGF) by the tumors. VEGFs act on the VEGF receptors (VEGFR) on endothelial cells, which orchestrates the angiogenesis cascade. Inhibition of angiogenesis has been an integral therapeutic approach in cancer. Multiple angiogenesis inhibitors have been developed and are currently in the clinic. **e.** Schematic depicting the consequence on the delivery of blood-borne nutrients to the tumors by angiogenesis inhibition. Apart from diminished oxygenation and elevation of hypoxia in the tumors, angiogenesis inhibition also restricts the import of blood-borne nutrients, that fuel the mitochondrial metabolic processes: the TCA cycle and oxidative phosphorylation. The TCA cycle is a bioenergetic engine and an anabolic hub in cancer cells. In addition to generating reducing equivalents for mitochondrial Oxphos reactions, TCA cycle also plays crucial role in anabolic reactions in cancer cells. The TCA cycle is constantly drained of carbon atoms in the form of CO_2_ and when the metabolic intermediates exit the cycle, to generate biosynthetic intermediates (shown in red) to support cancer cell proliferation. To ensure a continuous function of the TCA cycle, the carbon atoms must be replenished back into the TCA cycle by a process called anaplerosis. Cancer cells consume copious amounts of blood borne nutrients such as glucose, glutamine, fatty acids etc, to provide the surplus carbon atoms for the TCA cycle and ensure uncontrolled growth and proliferation.

**Supplemental Figure 3:**
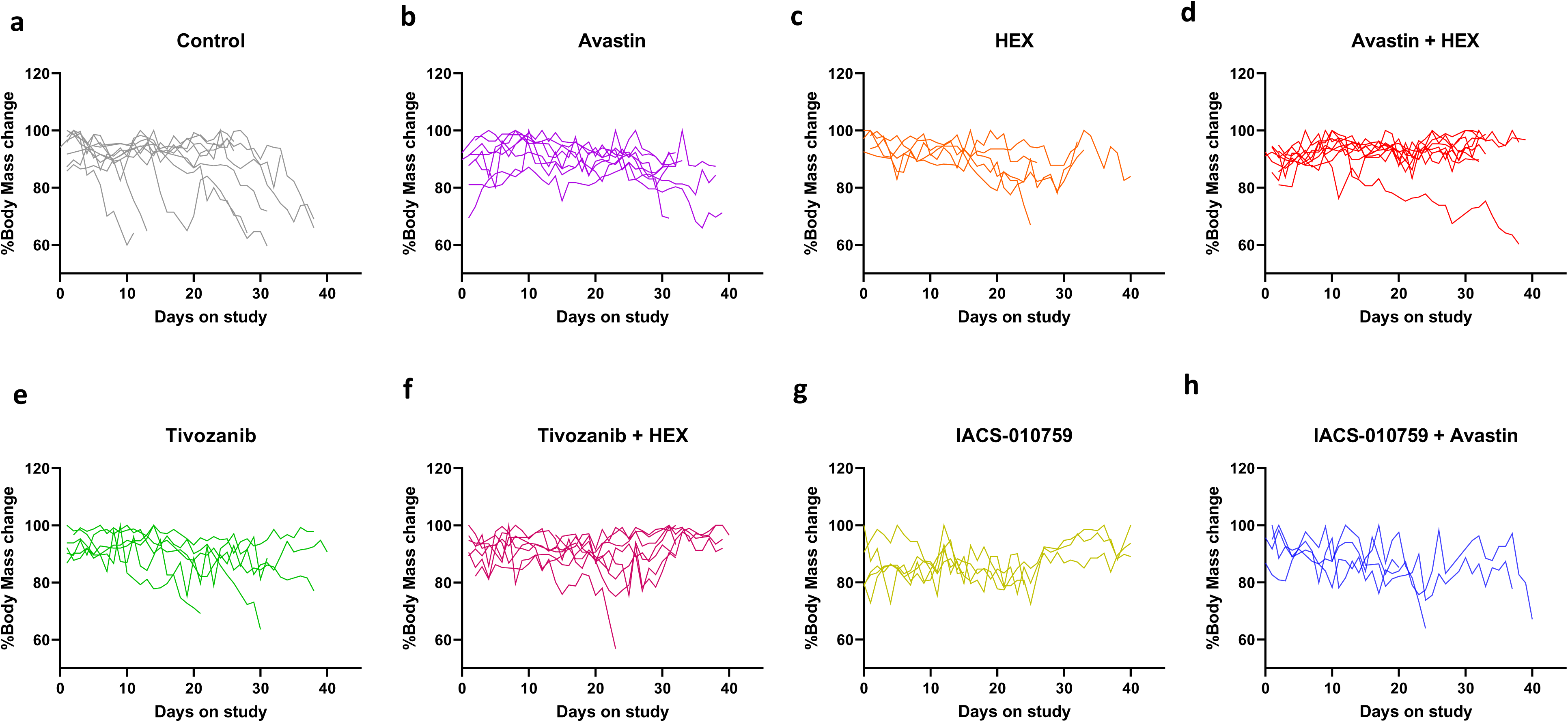
Inhibition of angiogenesis combined with anti-metabolic therapy is well-tolerated in-vivo. **a-e.** Body weight is a useful proxy of disease burden in mice that is routinely monitored in toxicology experiments. Body weights of mice bearing ENO1-/- intracranial tumors subjected to Avastin, HEX, Tivozanib or IACS-10759 as monotherapy or to combination of HEX or IACS-10759 with Avastin or Tivozanib over 28 days. Combination therapy does not exhibit toxicity exaggeration in mice compared to the monotherapy and delayed the loss of body weight caused by brain tumor disease burden. The most significant decreases in body weights occurred in mice in the control group and it correlated with most aggressive intracranial tumor growth. Body weights of individual mice show a negative correlation with tumor burden with loss of body weights strongly associated with rapid tumor growth.

**Supplemental Figure 4:**
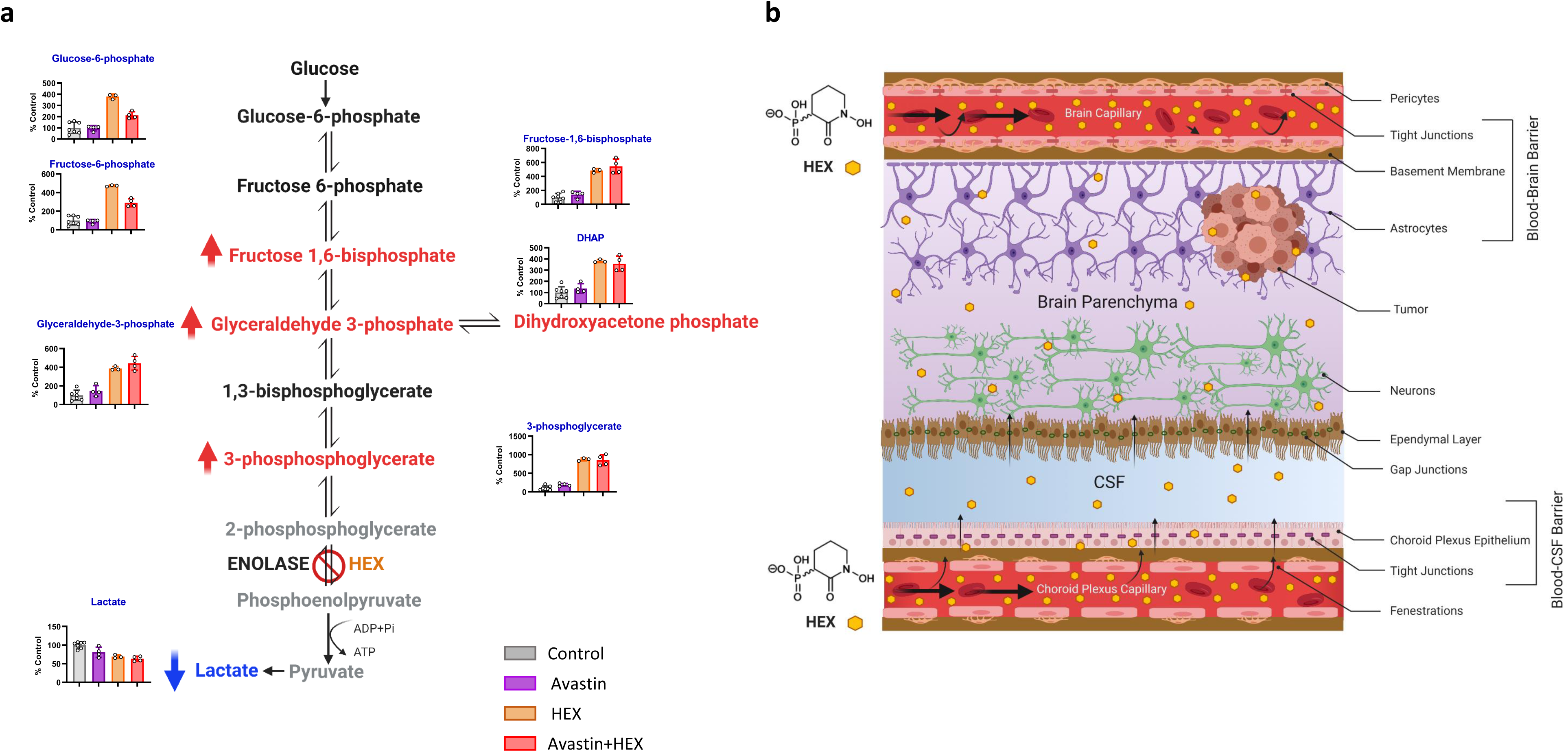
Avastin treatment does not modulate inhibition of glycolytic flux by the enolase inhibitor HEX. **a.** Inhibition of glycolysis by HEX is independent of Avastin mediated BBB resealing. HEX treatment, in HEX alone or Avastin + HEX treated tumors result in a comparable accumulation of glycolytic intermediates upstream of the enolase reaction. This indicates that despite the resealing of the blood brain barrier by Avastin treatment, HEX can still enter the brain and exert its effects. **b.** Small water-soluble phosphonate drug can reach the brain through the blood cerebrospinal fluid (CSF) barrier. Water-soluble phosphonate substrates such as HEX cannot cross the highly selective blood brain barrier (BBB). An alternative route for entry to the brain is through the blood CSF barrier. Unlike the BBB, the capillaries in the blood CSF barrier have fenestrations, which enable penetration of soluble drugs with small molecular weights, such as the antibiotic fosfomycin, which is clinically used to target brain abscesses. When Avastin treatment reseals the leaky tumor vasculature, HEX possibly enters the brain through the blood CSF barrier and exerts its anti-neoplastic effect (Lin *et al* 2020 *Nat Metab*, and Khadka *et al* 2020 *Cancer and Metab*)

**Supplemental Figure 5:**
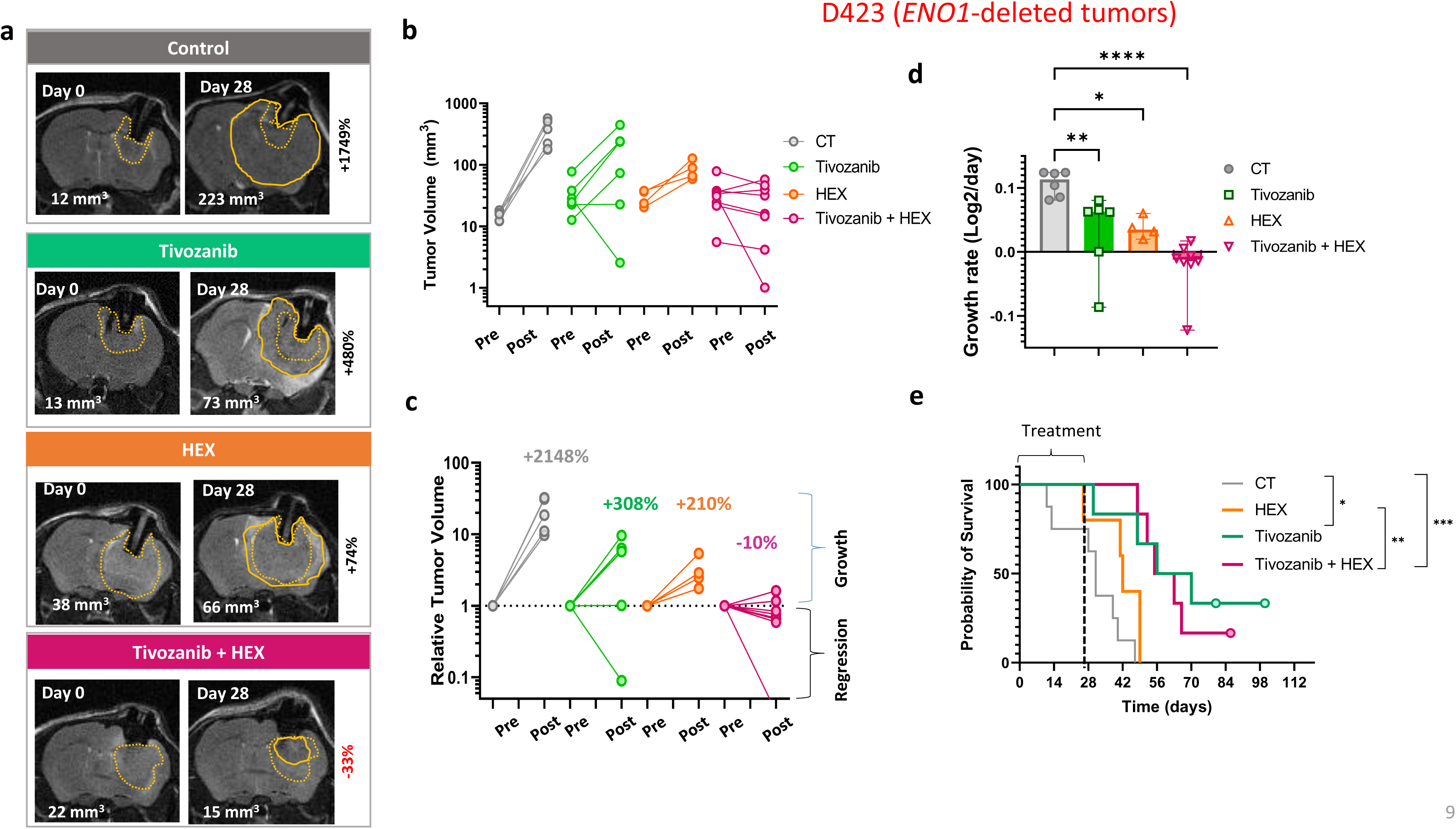
Combination of the enolase inhibitor HEX and VEGFR1/2/3 inhibitor Tivozanib all but eliminates intracranial tumor growth. Intracranial tumors were generated in Foxn1 *^nu/nu^* nude mice by implantation of D423 *ENO1*-homozygously deleted glioma cells and tumor growth was followed every week by T2-weighted MRI. When tumors were approximately 20 mm^3^ in volume, mice were separated into four groups: Control (n=6), Tivozanib (n=7), HEX (n=4), Tivozanib + HEX (n=8) and treatments with Tivozanib (2.5 mpk, 7X per week), Enolase inhibitor HEX (225 mpk, 14 doses per week) or Tivozanib plus HEX (Tivozanib, 2.5mpk 7X per week + 225 mpk SC 14X per week) were administered for 28 days. **a.** MRI images with tumor outlines before (dotted yellow) and after (solid yellow) 28 days of treatment. **b-d.** Treatment with Tivozanib as single agent, led to a modest inhibition of tumor growth in most mice, but in two mice, it completely eradicated the tumors. HEX as single agent attenuated tumor growth but did not result in an overall tumor regression. However, the combination of Tivozanib and HEX resulted in tumor regression in all treated animals. Animals were taken off the treatments on day 28 and survival in each group was calculated. Tivozanib as single agent as well as Tivozanib and HEX combination caused a significant extension of survival compared to control and HEX.

**Supplemental Figure 6:**
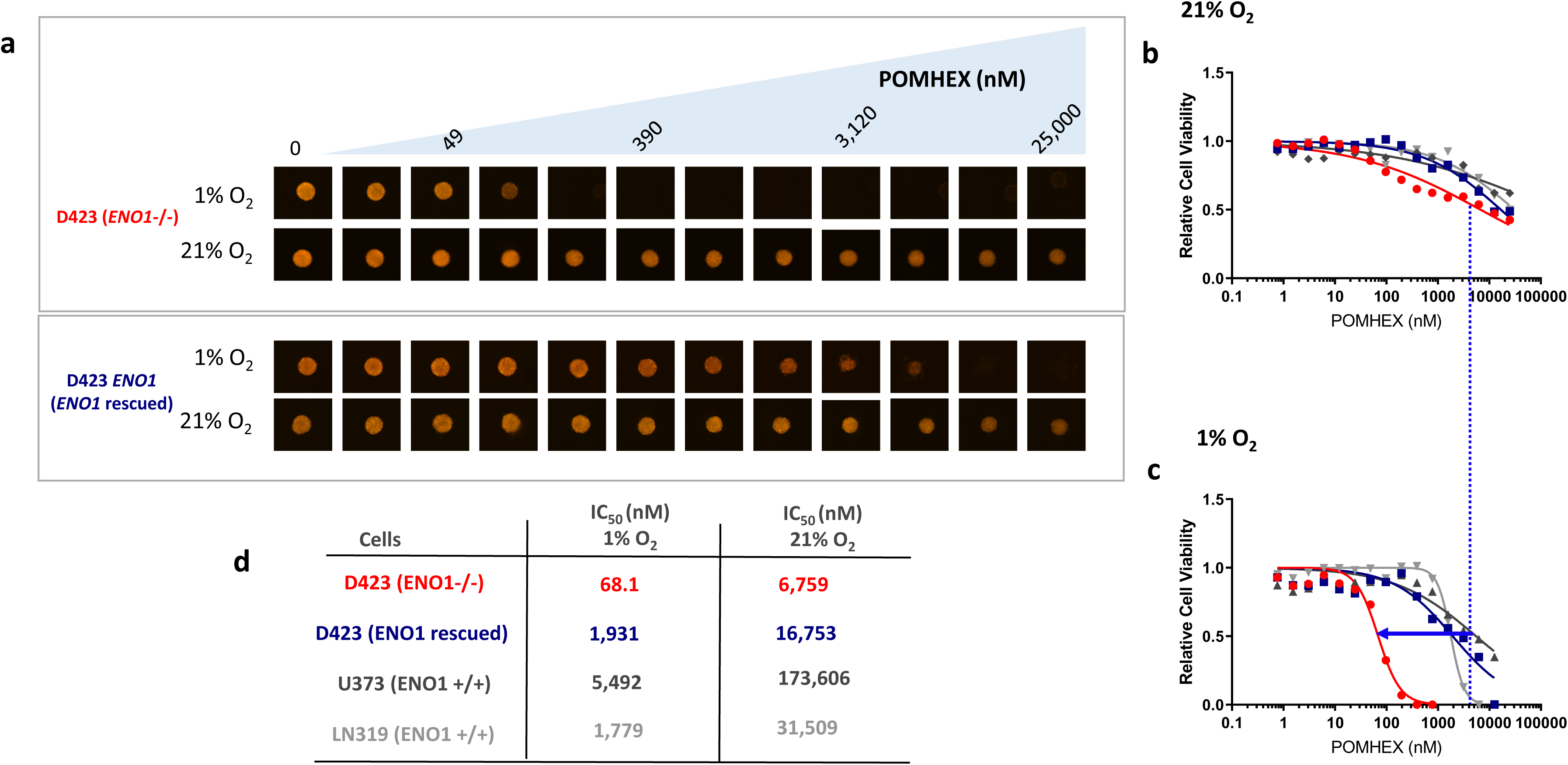
Potency of the Enolase inhibitor is dramatically enhanced under hypoxic conditions in 3D spheroids *in vitro*. Tumor spheres of D423 *ENO1*-deleted (red), D423 *ENO1*-rescued (blue), as well as *ENO1*-intact, U373 (dark grey) and LN319 (grey) cells were treated with POMHEX at the concentrations indicated, for 2 days in either a hypoxic incubator (1% O_2_) or under 21% O_2_ hypoxia. Viable cells were imaged by Tetramethylrhodamine (TMRE, red fluorescence) staining, which was quantified and expressed relative to the vehicle control. **a**. Representative images of *ENO1*-deleted (top panel) and *ENO1*-rescued tumor spheres (bottom panel) at normoxia or 1% hypoxia after 2 days of treatment with POMHEX at the concentrations indicated. **b-c.** Quantification of cell viability by TMRE intensity as a function of POMHEX concentration after two days of treatment under normoxia (**b**) or 1% hypoxia (**c**) for *ENO1*-deleted, *ENO1*-rescued and *ENO1*-WT lines treated with POMHEX. The dotted vertical line is for IC_50_ comparison, and the arrow indicates the shift in IC_50_ between hypoxic and normoxic conditions. **d.** Table indicating the IC**_50_**of each cell line under normoxic and 1% hypoxic conditions. The potency of the Enolase inhibitor POMHEX for killing D423 *ENO1*-deleted glioma cells is dramatically increased in 1% O_2_ as compared to in normoxia; hypoxia similarly increases potency of POMHEX against *ENO1*-rescued and *ENO1*-WT glioma cell lines.

**Supplemental Figure 7:**
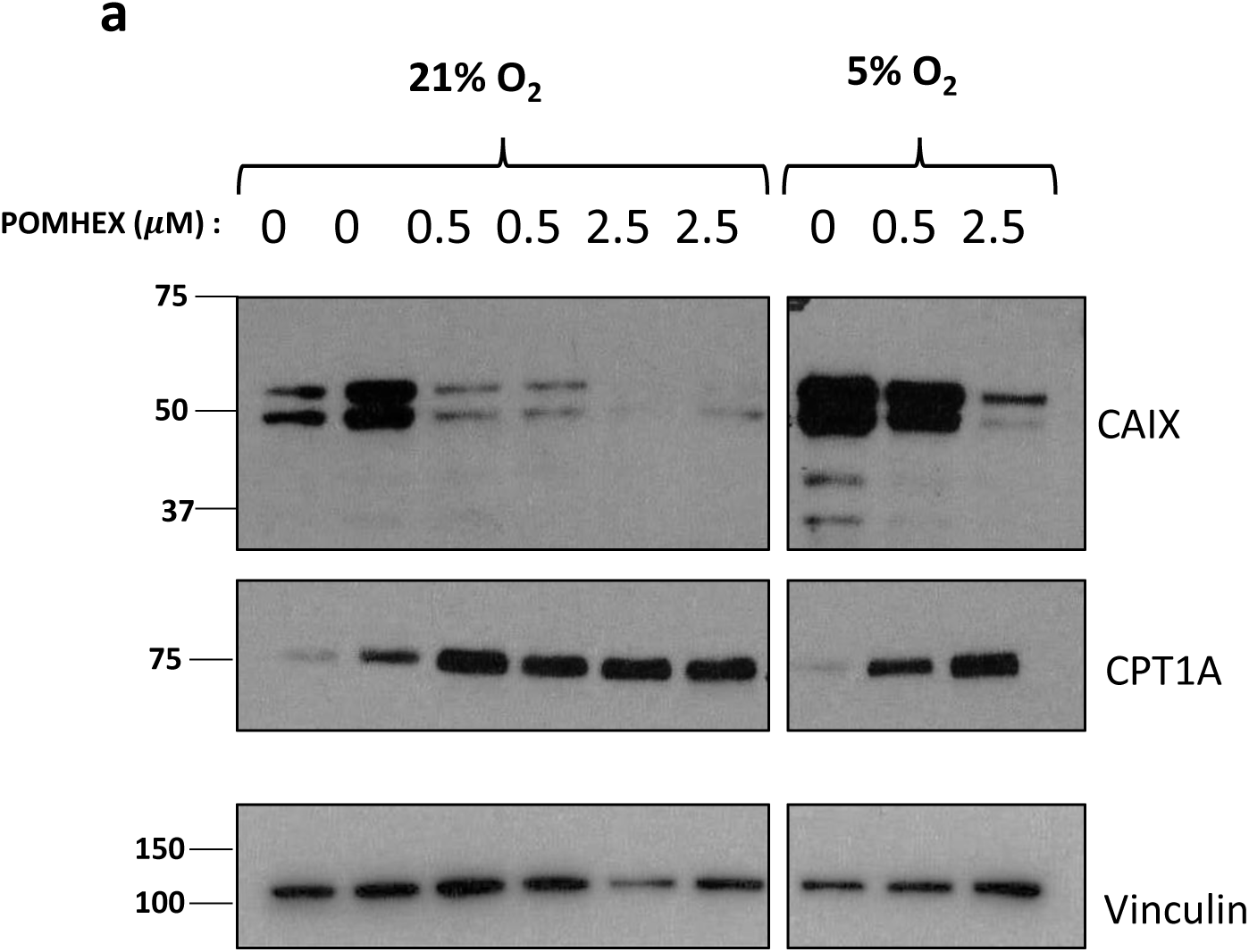
Enolase inhibitor treatment decreases hypoxia by diminishing oxygen consumption. Enolase inhibitor POMHEX inhibits glycolysis and prevents pyruvate production, a key mitochondrial oxidative phosphorylation (oxygen consuming) substrate. Cells were treated different doses of POMHEX in normoxic and hypoxic conditions and different hypoxia responsive genes were assessed by western blot. **a.** Treatment with the Enolase inhibitor decreases carbonic anhydrase-9, a HIF1α target in both normoxic and hypoxic conditions in a dose dependent manner but increases carnitine palmitoyl-transferase CPT1A levels.

**Supplemental Figure 8:**
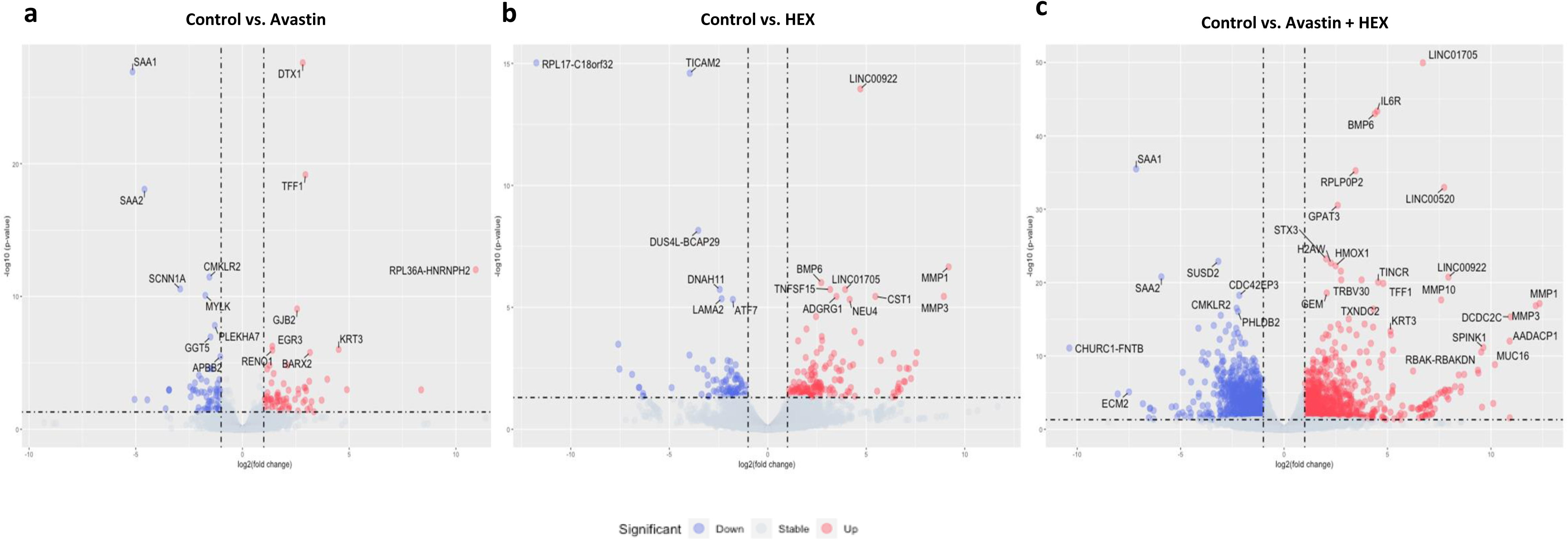
Combination of Avastin and HEX causes synergistic changes in differential gene expression in intracranial tumors. (**a-c**) Volcano plots representing differentially up-regulated or down-regulated genes in monotherapy of Avastin or HEX, and combination treatment with Avastin and HEX compared to control. Statistically significant up- (red)or down- (blue)regulated genes (Up: log2Fc ≥ 1 and padj ≤ 0.05; Down: log2Fc ≤ −1 and padj ≤ 0.05) are shown in red and blue circles respectively, with gene names highlighted.

**Supplemental Figure 9:**
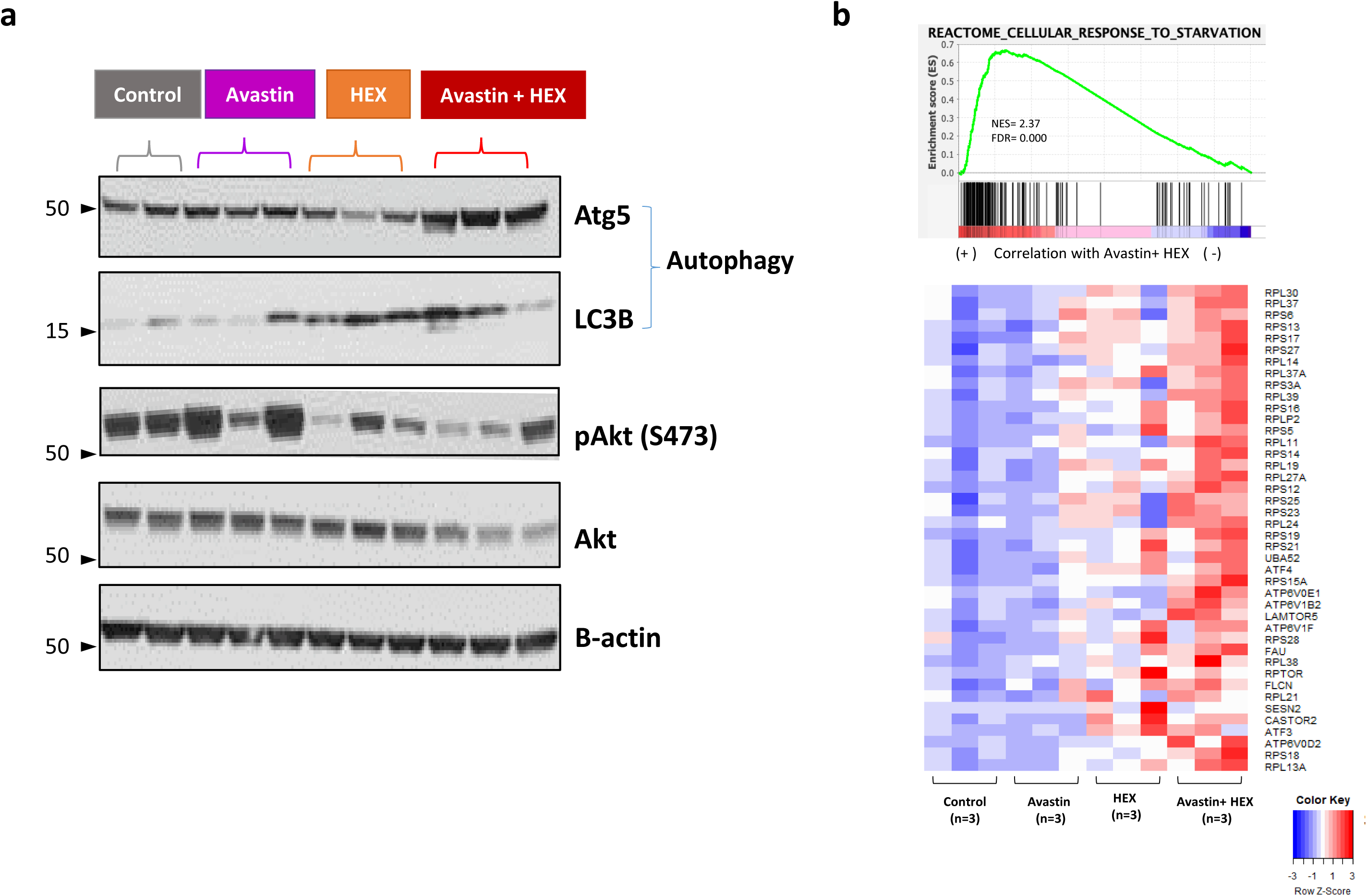
Combination of angiogenesis and glycolysis inhibition accentuate nutrient stress and adaptation response in tumors. **a.** Lysates from intracranial tumors from control, Avastin, HEX, and Avastin + HEX treatment groups were immunoblotted for proteins such as Atg5 and LC3B involved in autophagy mediated stress adaptation response, and Akt, a crucial protein at the intersection of many interconnected signaling pathways that control cell survival and growth, apoptosis, as well as cellular metabolism. Combination treatment of Avastin and HEX significantly accentuates autophagic response which is evidenced by increase in Atg5 as well as LC3B compared to monotherapy of HEX or Avastin, and also suppress Akt signaling, by reducing levels of Akt protein as well as its phosphorylation. **b**. GSEA plots showing positive enrichment of genes in the reactome cellular response to starvation gene set in Avastin + HEX group compared to control (**Also shown in** Figure 4i). **c.** The leading-edge genes for all four treatment groups are shown in the heatmap.

**Supplemental Figure 10:**
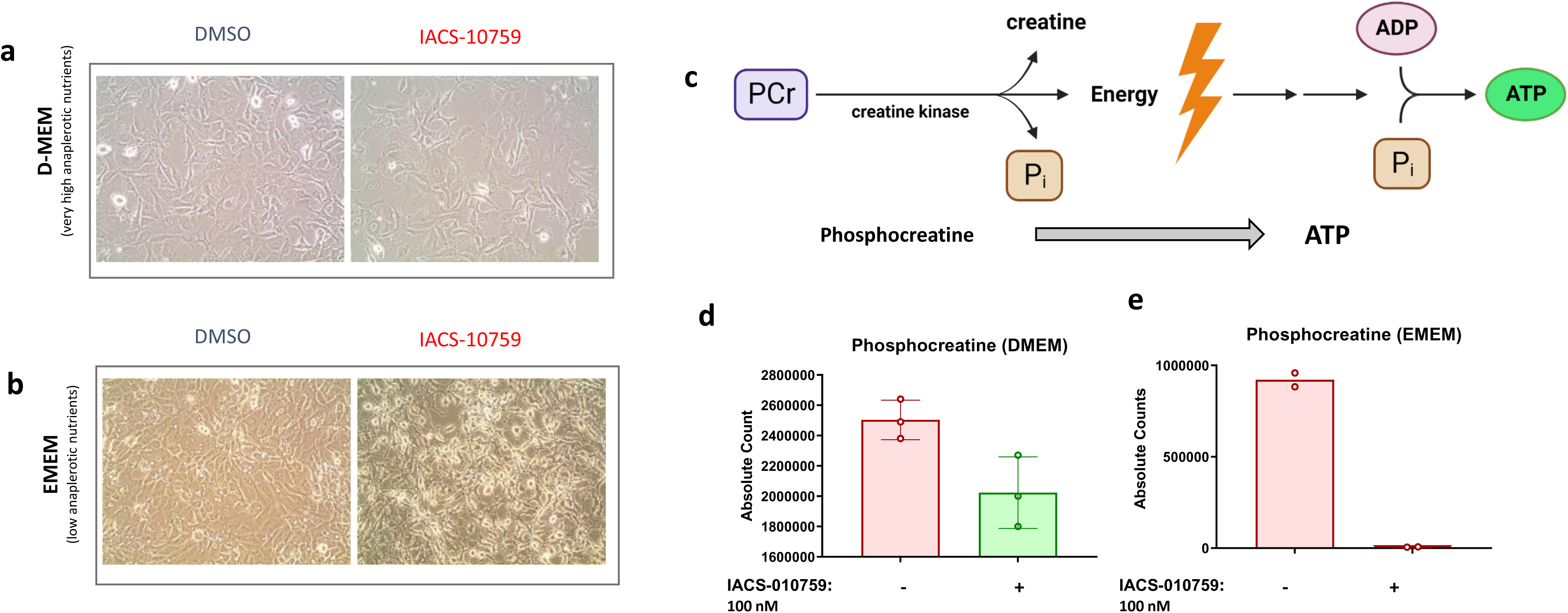
Inhibition of oxidative phosphorylation induces anaplerotic stress, resulting in an exaggerated bioenergetic collapse in nutrient deficient conditions. **a-b**. *ENO1* deleted D423 cells treated with IACS-10759 in nutrient replete (**a**) and nutrient depleted conditions (**b**) show that the toxicity of IACS-010759 is exaggerated in low nutrient conditions. **c-e.** Phospho-creatine level, an indicator of bioenergetic state of cells (**c**), reveal a profound disruption of bioenergetics by IACS-010759 treatment in nutrient deficient conditions (**e**) compared to nutrient rich condition (**d**).

**Supplemental Figure 11A and B:**
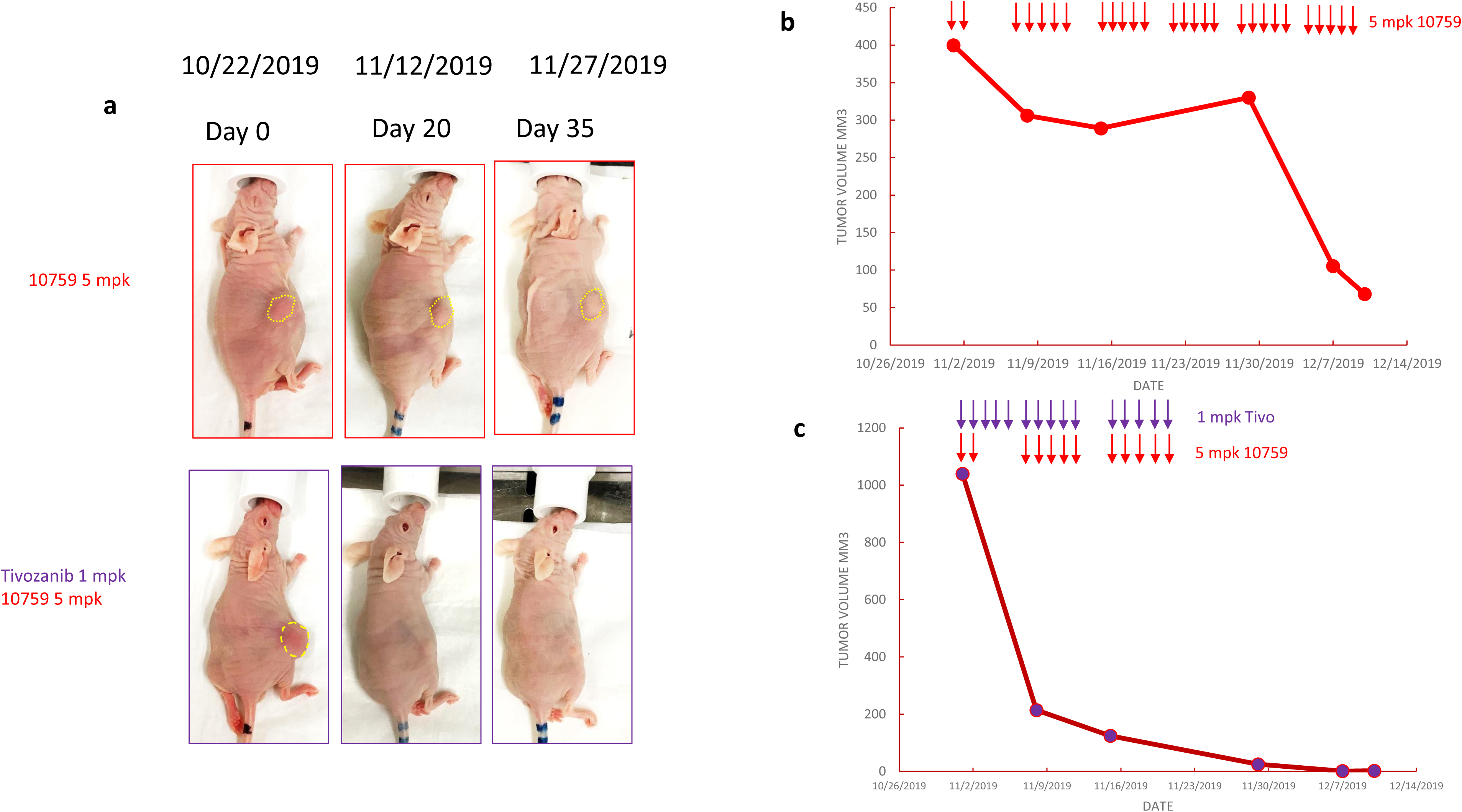
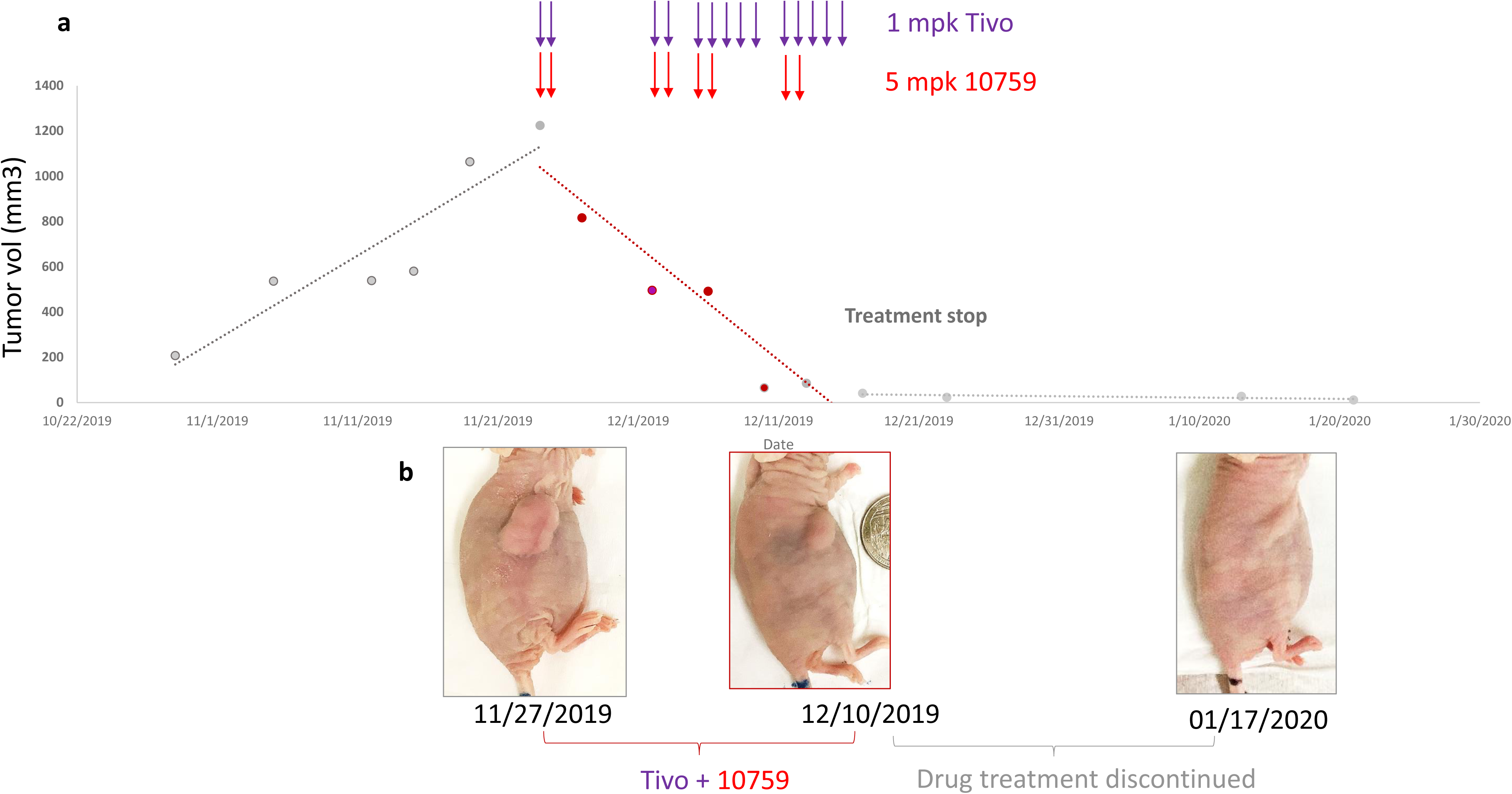
Angiogenesis inhibitor and IACS-010759 drive exceptionally rapid regression even in very large sub-cutaneous tumors. *PGD-*homozygously deleted NB1 cells were implanted sub-cutaneous in immunodeficient Foxn1 Nude mice. **12A a-c.** When tumors reached large volumes (>400 mm^3^) mice were treated with either IACS-010759 alone (each dose indicated by a red arrow) alone or with IACS-010759 plus Tivozanib (each dose indicated by a purple arrow). Tivozanib as a monotherapy decreased tumor growth while IACS-010759 abolished tumor growth and as reported previously – drove tumor regression slowly over the course of weeks of continuous treatment. However – the co-administration of IACS-010759 with Tivozanib led to massive regression (>75% reduction in volume) in only 5 days. **12B a-b.** Given the poor tolerability of IACS-010759 in the clinic; dose-reduction attempts were made here: it was found that only two doses of IACS-010759 per week along with 5 doses of Tivozanib were sufficient to drive complete tumor regression with tumors remaining dormant for extended time after treatment discontinuation. Further experiments were discontinued due to COVID19 shut-downs.

**Supplemental Figure 12:**
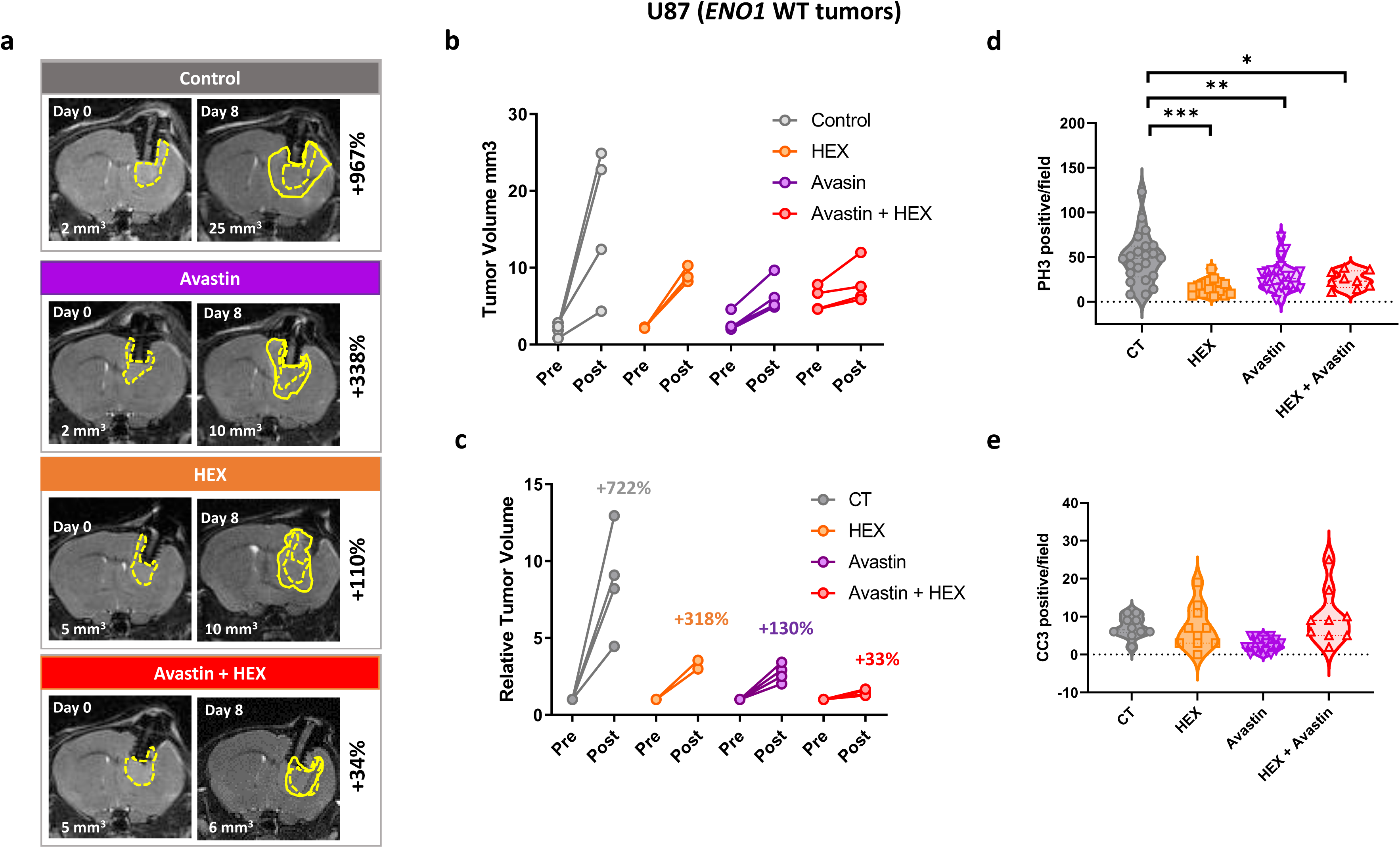
Angiogenesis inhibitor and enolase inhibitor display synergistic activity against non-glycolysis-compromised tumors. Intracranial tumors were generated by implanting U87 *ENO1* intact glioma cells in NSG immunocompromised mice. Tumor development was followed by T2-MRI. Treatment was begun when tumors reached ~2 mm3. **a.** T2-weighted MRI images of animals before and after 8 days of treatment with tumor volumes indicated in mm^3^ in the lower part of the image; initial tumor outlines are shown in dotted yellow lines, while tumors after 8 days are shown in solid lines. **b-c.** Summary of tumor volume changes after 8 days on treatment. Animals were treated continuously with a high dose of HEX, once daily for 8 days, while Avastin was administered at 200 mg/kg 3 X in 8 days. The effect of HEX in ENO1-WT gliomas is only marginal, but Avastin treatment led to a modest inhibition of tumor growth. Combination of HEX and Avastin result in a near-all suppression of tumor growth. **d-e.** Histopathological analyses brain sections extracted from mice showed a significant reduction in (phospho-histone H3 positive, an index of proliferation) in tumors treated with the combination of HEX, Avastin and Avastin and HEX, and an increase in dying cells (cleaved caspase 3 positive cells, an index of apoptotic cells), in Avastin and HEX treated tumors.

**Supplemental Figure 13:**
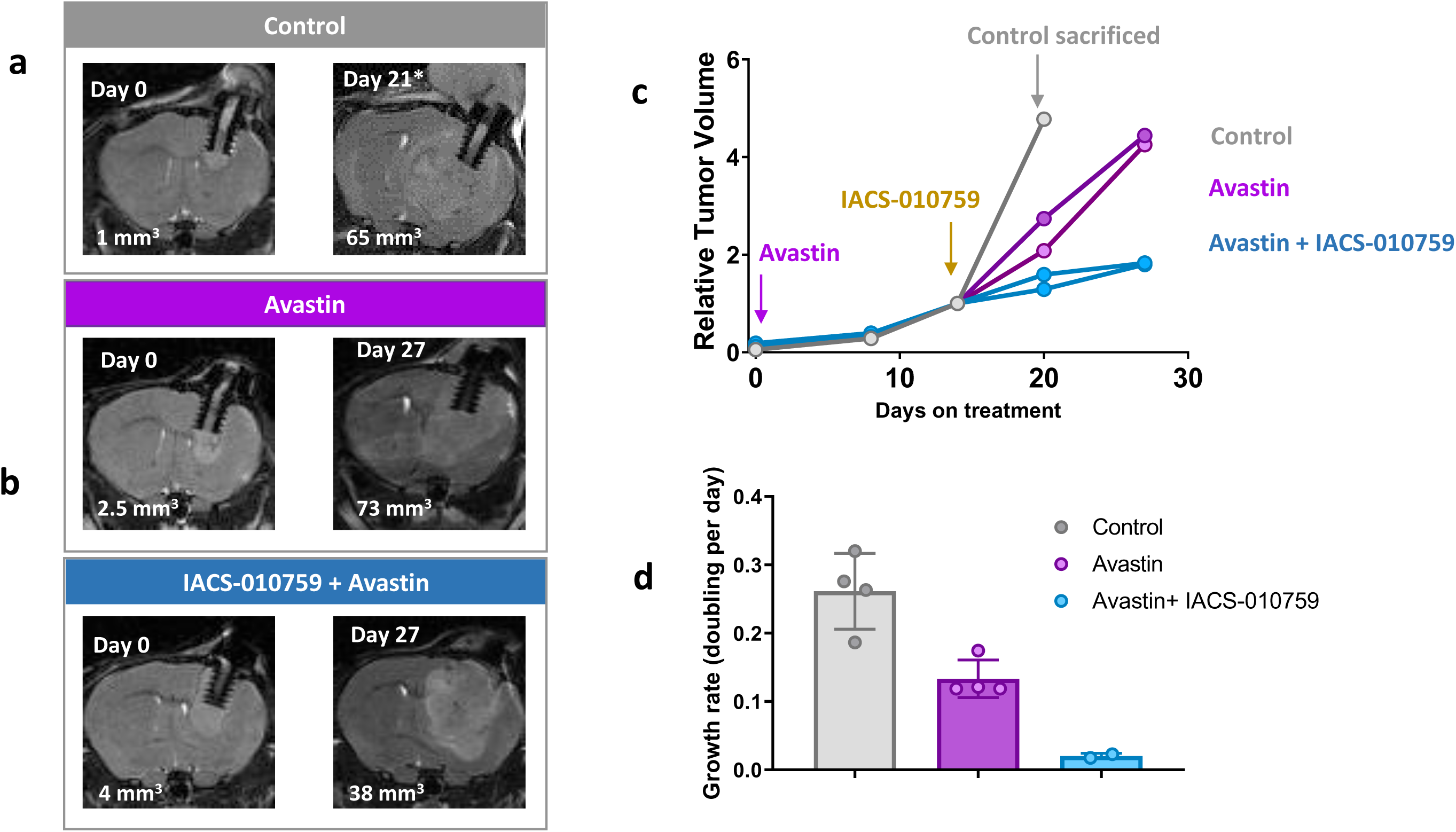
Angiogenesis inhibitor and IACS-010759 display synergistic activity against non-glycolysis-compromised tumors. *ENO1* WT U87 cells were implanted intracranially in immunodeficient NSG mice. Tumor bearing mice were treated with Avastin alone or with combination of Avastin and IACS-010759. **a-d.** Tumor volumes pre-treatment are outlined with dotted yellow lines (Day0) and post-treatment are outlined with solid yellow lines (Day 27) **(a-b)** Avastin as a single agent moderately delays tumor growth, but Avastin strongly sensitizes *ENO1* intact tumors to inhibition of mitochondrial oxidative phosphorylation, resulting in stasis of tumor growth.

**Supplemental Figure 14:**
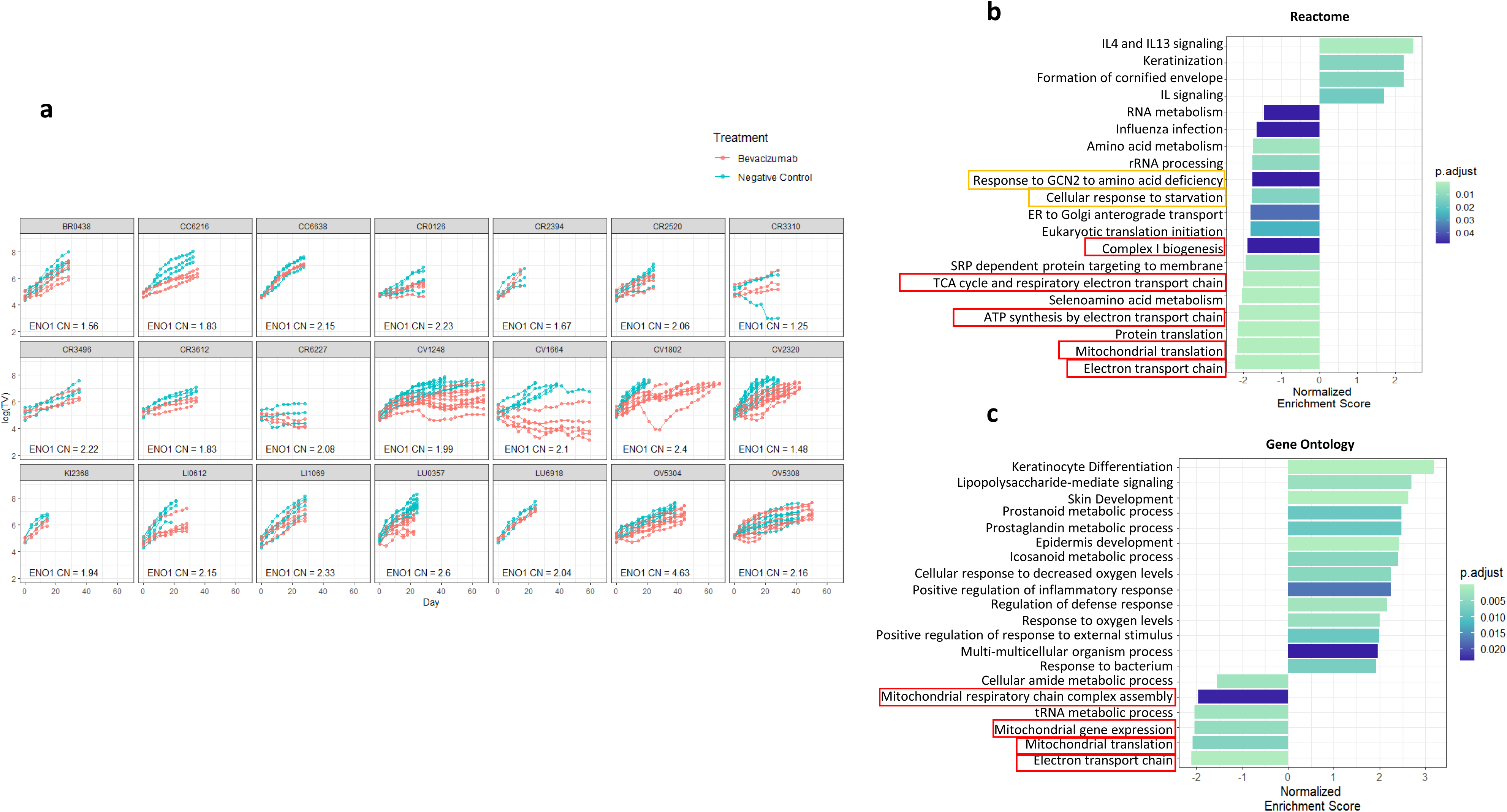
Low expression mitochondrial OxPhos gene is associated with favorable response to Avastin in a broad panel of PDXs. PDXs in the CrownBio collection differing in anti-tumor responsiveness to Avastin (For example: CV1664 highly responsive vs CV2320 minimally responsive) none of which is *ENO1*-homozygous deleted (ENO1 CNV = 0) **a.** Relative tumor volume versus time on treatment shown for individual mice treated with vehicle group (green) and Avastin(red) dosed at 10 mg/kg IP once per week. **b-c**. The tumors were profiled (pre-treatment) by RNA seq and human transcript reads (genes expressed in malignant cancer cells) were analyzed for markers predicting response to Avastin. Tumor growth data was combined with RNAseq data to determine the effect of Avastin treatment on tumor growth as well as gene:avastin interaction for each gene. GSEA analyses and normalized enrichment scores for reactome (**b**) and gene ontology (**c**) datasets are shown. For GSEA, the significant human genes (p.adj < 0.05, n = 5419) were ranked by 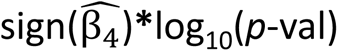. A positive NES indicates that activation of the pathway synergizes with Avastin treatment, whereas a negative NES indicates an antagonistic effect. Key findings indicate that those PDXs with lower expression of TCA-cycle and mitochondrial genes (red box) showed a better response to Avastin, while those with higher expression of these genes antagonized anti-tumor effect of Avastin. Similarly, low expression of the amino acid starvation response (orange boxes) was also associated with favorable response to Avastin.

